# Oxygen metabolism in descendants of the archaeal-eukaryotic ancestor

**DOI:** 10.1101/2024.07.04.601786

**Authors:** Kathryn E. Appler, James P. Lingford, Xianzhe Gong, Kassiani Panagiotou, Pedro Leão, Marguerite Langwig, Chris Greening, Thijs J. G. Ettema, Valerie De Anda, Brett J. Baker

## Abstract

Asgard archaea were pivotal in the origin of complex cellular life. Hodarchaeales (Asgardarchaeota class Heimdallarchaeia) were recently shown to be the closest relatives of eukaryotes. However, limited sampling of these archaea constrains our understanding of their ecology and evolution^1–3^, including their anticipated role in eukaryogenesis. Here, we nearly double the number of Asgardarchaeota metagenome-assembled genomes (MAGs) to 869, including 136 new Heimdallarchaeia (49 Hodarchaeales) and several novel lineages. Examining global distribution revealed Hodarcheales are primarily found in coastal marine sediments. Detailed analysis of their metabolic capabilities revealed guilds of Heimdallarchaeia are distinct from other Asgardarchaeota. These archaea encode hallmarks of aerobic eukaryotes, including electron transport chain complexes (III and IV), biosynthesis of heme, and response to reactive oxygen species (ROS). The predicted structural architecture of Heimdallarchaeia membrane-bound hydrogenases includes additional Complex-I-like subunits potentially increasing the proton motive force and ATP synthesis. Heimdallarchaeia genomes encode CoxD, which regulates the electron transport chain (ETC) in eukaryotes. Thus, key hallmarks for aerobic respiration may have been present in the Asgard-eukaryotic ancestor. Moreover, we found that Heimdallarchaeia is present in a variety of oxic marine environments. This expanded diversity reveals these Archaea likely conferred energetic advantages during early stages of eukaryogenesis, fueling cellular complexity.

## Main

The discovery of Asgard archaea (Asgardarchaeota phylum) has revolutionized our understanding of the origins of complex life and the shape of the tree of life^1,4^. These microbes are the descendants of the host that gave rise to eukaryotes, containing eukaryotic-like proteins^5,6^. Models of eukaryogenesis predict the archaeal host to be an anaerobe (e.g., HH^7^, HM/HS Syntrophy^8,9^, and Entangle-Engulf-Endogenize^10^) or facultative anaerobe/aerobe (e.g., Reverse Flow^2^) that conserved energy via substrate-level phosphorylation through fermentation of organic compounds to hydrogen and acetate. Recently, it was predicted the first eukaryotes diverged from Hodarchaeales, a heterotrophic lineage adapted to mesophilic environments^1^, and Alphaproteobacteria about 2.67-2.19 billion years ago (Ga) and 2.58-2.12 Ga, respectively^11^ (earlier than previous reports <1.84 Ga)^12^. Before this, there was a rise in oxygen on the planet ∼2.5-2.1 Ga, referred to as the Great Oxidation Event (GOE)^13,14^. Most eukaryotes today are aerobes, while the anaerobes have lost their ability to use oxygen^15^. It is plausible that the last archaeal and eukaryotic common ancestor (LAECA) and potentially their modern descendants, Hodarchaeales, could use oxygen. Yet, most contemporary Asgard archaea are inferred to be strict anaerobes recovered from anoxic and microoxic environments,^2,10,16^ though a few reports suggest some Heimdallarchaeia are capable of aerobic or nitrate respiration^2,16,17^.

Current eukaryogenesis models are based on the metabolic capabilities of a few extant representatives of Asgard, including two cultured representatives of Lokiarchaeales^10,18^. However, these lineages are very distinct from Hodarchaeales (Heimdall LC-3), proposed to be the closest prokaryotic relatives of eukaryotes^1,2^. The limited genomic sampling of these lineages, currently just 23 MAGs available, has prevented the detailed reconstruction of the metabolic capabilities of LAECA. The most recent report of these lineages with a limited number of genomes suggests they are anaerobic respiratory microorganisms, using organic carbon and hydrogen as energy sources and nitrate as their electron acceptor^2,16,17^. However, an in-depth examination of the metabolic capacity of the lineages most closely related to eukaryotes, the Heimdallarchaeia, has thus far been lacking.

Here, we conducted a massive DNA sequencing of coastal and deep-sea sediments to expand the genomic diversity of Asgard archaea, nearly doubling their known diversity and providing the first detailed metabolic description. This expanded catalog enabled us to explore the metabolic and bioenergetic potential of Hodarchaeales to advance our understanding of their roles in the origin of eukaryotes. Our analysis revealed that lineages most closely related to eukaryotes are capable of high energy-yielding aerobic metabolism.

### Expansion of Asgard genomic diversity

To expand the genomic catalog of these archaea, we chose two ocean floor ecosystems rich in Asgard diversity, the shallow (17-30.5 m) coastal Bohai Sea (BH, Eastern China) and the deep sea hydrothermal system in (2,000 m) Guaymas Basin (GB, Gulf of California). We combined 10 Tb of new sequencing (GE3; the largest known deep-sea genomic database) with previously reported efforts (BH & GE2)^19–21^ and extensive manual curation to obtain over 13,000 MAGs (see methods). From these, a total of 404 MAGs were classified as Asgardarchaeota, expanding the known diversity of this group by 46.5% (Fig. 1). We also identified several Asgard representatives mislabeled as other archaea in public databases.

**Fig. 1.**
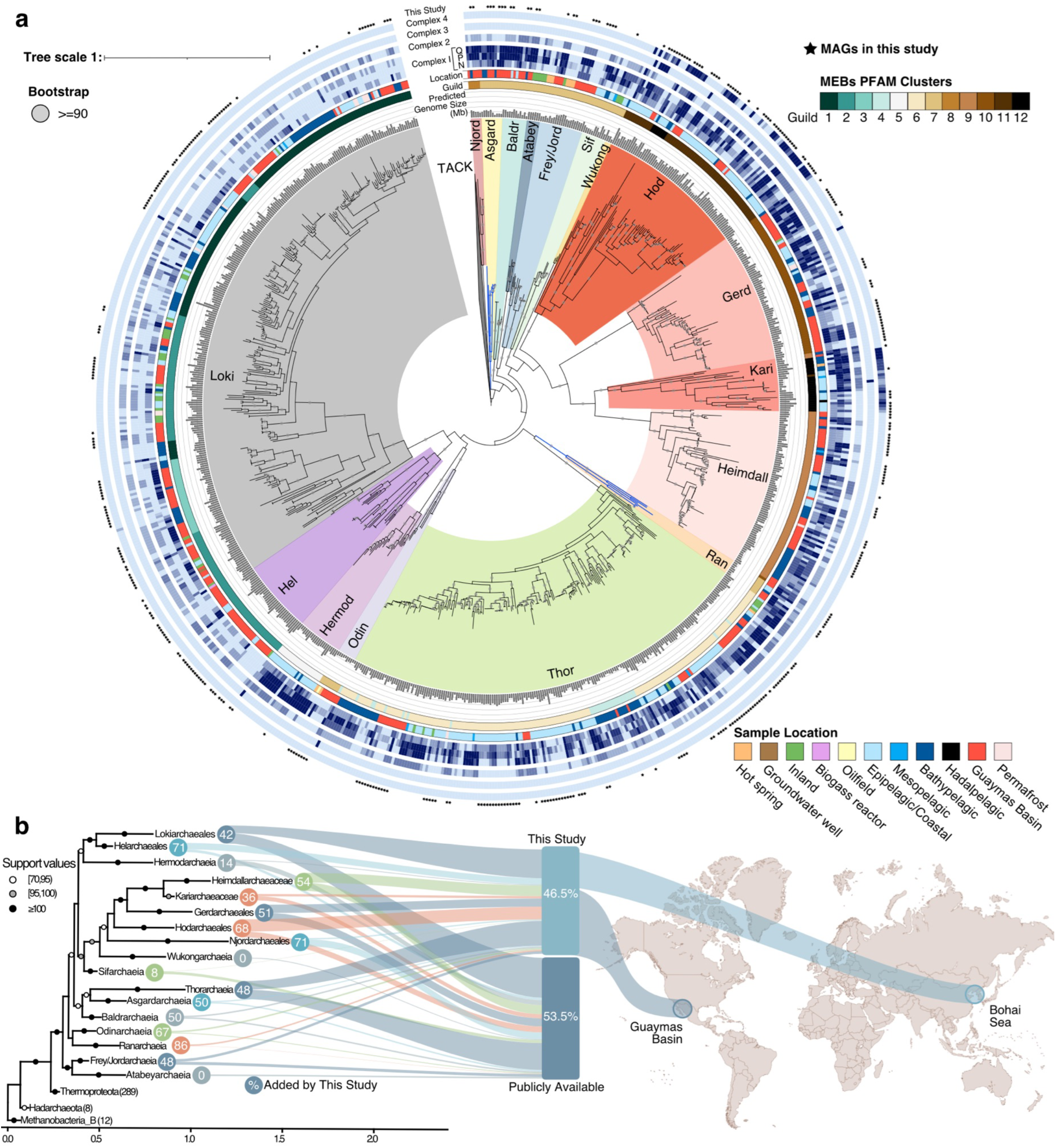
| Expanded diversity of Asgard archaea. **a**. Maximum likelihood phylogeny based on 47 arCOGs for the 837 Asgard MAGs and 36 TACK archaea representatives as the outgroup. The tree was generated using RAxML v8.2.11 with free model parameters estimated by the RAxML GAMMA model of rate heterogeneity and ML estimate of alpha-parameter–bootstraps based on 1000 rapid bootstrap inferences. The blue branches (lower right) indicate the new Asgararchaeota class, Ranarchaeia and the recently proposed Asgardarchaeia (top). The concentric rings (in-out) highlight the predicted genome size (Mb), metabolic guilds based on Pfam clustering, sampling locations, subunits in the electron transport chain Complex I-IV (Complex I modules are shown separately), and black stars for the MAGs added by this study. **b**. Mapped on the phylogeny, using the new NM47 markers (see methods), is the percentage of MAGs from each lineage added by this study (circle), including 46.5% of the catalog from the Guaymas Basin (65.3%) and the Bohai Sea (34.7%). These MAGs constitute a 115.1% increase in the medium-high quality publicly available genomes.

To determine the identity of these new MAGs, we employed robust phylogenetic methods using a variety of marker sets (16S rRNA genes, 47 arCOG, 37 Phylosift, NM47^22^ and evolutionary models^1,23,24^. We recovered the previously suggested monophyly of Heimdallarchaeia when saturation and compositional bias were addressed (Fig1b and see methods)^1^. The 404 new Asgard archaea MAGs span 13 known lineages (Loki-, Hel-, Hermod-, Thor-, Frey/Jord-, Odin-, Baldr-, Sif-, Heimdall-, Kari-, Gerd-, Njord-, and Hodarchaeaceae), one new class Ranarchaeia (see taxonomic description below), and one metabolically uncharacterized class (Asgardarchaeia)^25^. Ranarchaeia is a deeply-branching Asgardarchaeota lineage (in the NM47 phylogenies). This expanded genomic diversity constitutes a valuable resource, increasing support in the phylogeny for the deep-rooted clades and improving the resolution at lower taxonomic ranks, including 49, 56, and 178 MAGs with unnamed orders, families, and genera (based on GTDB-tk v2.1.1), respectively^26^. For example, a parallel study by Köstlbacher et al., identified novel eukaryotic signature proteins (ESPs) via structural predictions in these MAGs (Köstlbacher et al., in prep). This genomic catalog substantially enriched the diversity of evolutionarily important Asgard groups, with an increase of 68% of the Hodarchaeales representatives, the closest known ancestor to eukaryotes.

### Hodarchaeales and Kariarchaeaceae are adapted to aerobic environments

To better understand the modern distribution of these archaea, we compared the environmental conditions from which the 869 Asgard MAGs were obtained, which fell into 12 categories (Fig. 2). Identification of previously uncharacterized Asgard archaea from public databases and curation of metadata allowed us to expand the known ecological ranges of these organisms. We find that Asgardarchaeota is present in a wide range of environments, from hot (91°C; hot springs^5,27–29^ and hydrothermal vents^1,3,5,6,17,30–33)^ to cold (∼-7°C; permafrost^34^), surface (<200 m deep^17,35–37)^ to deep (>6000 m^38^) oceans, freshwater to hypersaline (116.6 ppt^39^), shallow to the deep sediments (280.17 mbsf^40^), and in human-made structures (e.g., biogas reactors^41^). Despite most clades spanning ecological niches, Atabeyarchaeia seems unique to wetland soils and inland lakes^40,42^, and Njordarchaeales and Odinarchaeia are exclusively associated with hydrothermal-associated locations. Lokiarchaeia and Thorarchaeia have the greatest number of MAGs and are the most widespread^5,30,43^.

**Fig. 2.**
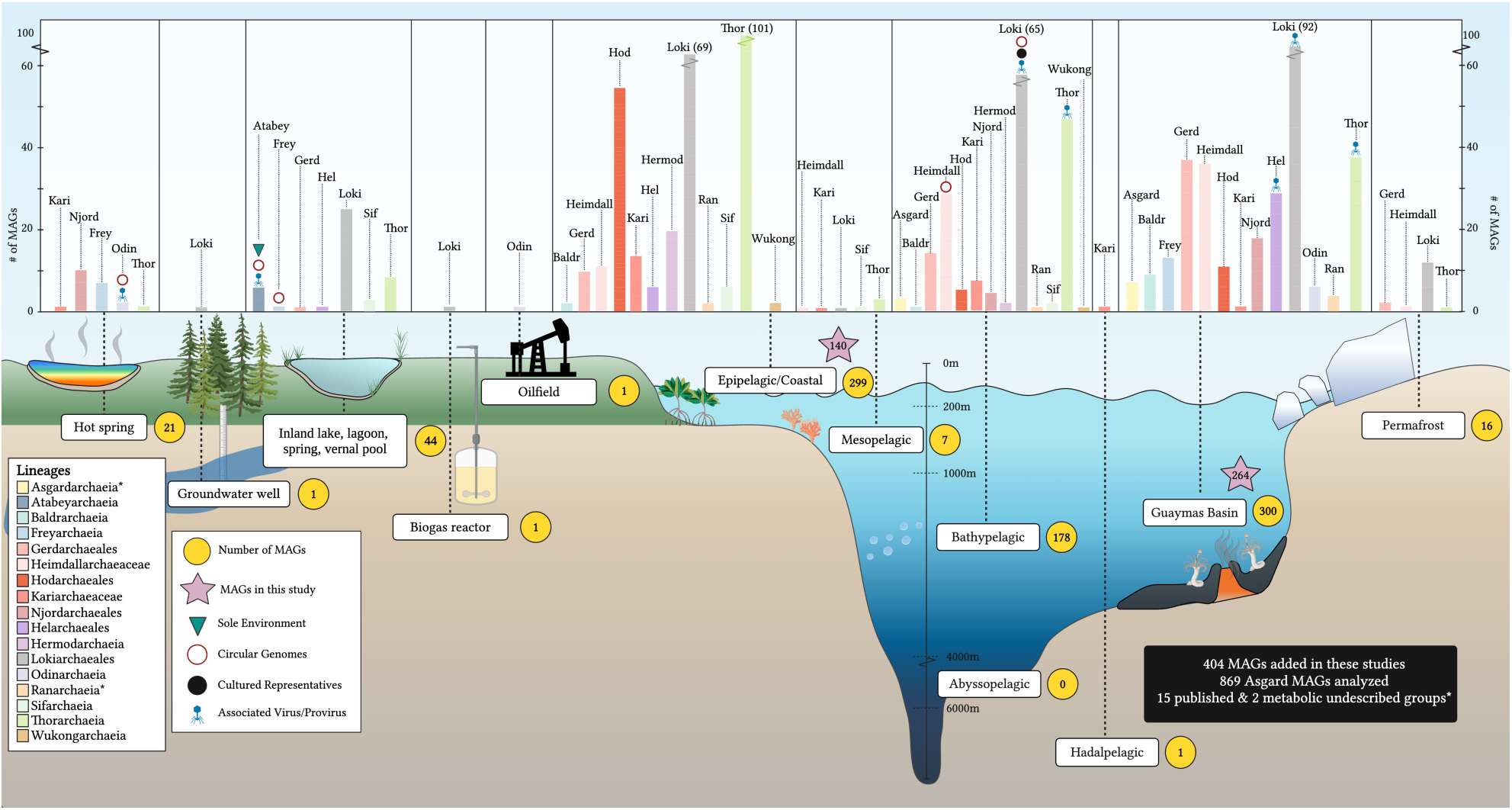
| Distribution of Asgardarchaeota based on the recovery of MAGs from a wide variety of habitats. The ecological distribution of the Asgardarchaeota genomes across 12 environments, including genomes added by this study (purple star) and the total number of MAGs (yellow circle). The bar chart shows the number of MAGs from each lineage by environment, indicating lineages found only in a single environment (Atabeyarchaeia; teal triangle), circular complete genomes (black circle), and MAGs with known associated viruses or proviruses (blue phage). We note the novel class, Ranarchaeia, and recently described Asgardarchaeia in the legend with asterisks. Guaymas Basin is shown separately from the other bathypelagic-associated MAGs to showcase the increased genomic diversity from this site.

Within the five Heimdallarchaeia clades, we identified potential niche partitioning. Most Heimdallarchaeaceae, Gerdarchaeales, and Njordarchaeales MAGs were extracted from the bathypelagic (1000-4000 mbsl). Yet, thus far, Hodarchaeales and Kariarchaeaceae have been most commonly associated with epipelagic sediments (<200 mbsl), including 78% and 56% of the MAGs from each lineage. To investigate potential aerotolerance in Heimdallarchaeia, we identified six of the shallow Kariarchaeaceae recovered from likely aerobic environments, including the Chattahoochee River (USA), Geoffrey Bay (Australia), Persian Gulf (Iran) (DO: 7.2 mg/L), and the Red Sea (Saudi Arabia) water column^35–37,44^. Most Asgard-focused studies have not reported oxygen levels because the lineages were predicted to be anoxic. In lieu of publicly reported oxygen concentrations, we used modeling based on amino acid frequencies. Of all the 869 Asgardarchaeota representatives, only 22 Kariarchaeaceae and 32 Hodarchaeales MAGs were predicted to be obligate or facultative aerobes^45^.

The mitochondrial ancestor is predicted to have been a facultative aerobic Alphaproteobacteria-related lineage^46,47^. Within the Heimdallarchaeia class, Hodarchaeales had the most MAGs co-occurring with Alphaproteobacteria (20 orders), with all but one MAG from the coastal Bohai Sea sediments, providing the first evidence of co-occurrence between these lineages across sites and depths. Aerobic Kiloniellales had the most co-occurring MAGs, yet we identified several unnamed lineages. Additionally, Rhodobacterales and Sphingomonadales (aerobes), as well as Rhizobales (aerobe/facultative anaerobe) and Rickettsiales (debated relation with mitochondrial ancestor), are also found in the same depth horizons as Hodarachaeales. We searched public genomic databases and identified 99.4% (824/829) and 100% (998/998) of Hodarchaeales and Kariarchaeaceae samples, respectively, which also contain Alphaproteobacteria^48,49^. Within these samples, we identified known aerobic lineages Acetobacter, Caulobacterales, Kiloniellales, and Sphingomonadales in 453, 626, 604, and 666 samples, respectively, co-occurring with Hodarchaeales.

### Evolutionary and ecological transitions in metabolic traits

To distinguish lineage and environmental-specific metabolic features, we clustered the genomes into defined groups based on their shared protein composition (see methods), as previously described^20^. The 12 Asgardarchaeota guilds have comparable metabolic functions (Fig. 1a). We also used pangenomic analyses to identify a set of uniquely conserved protein-coding genes for each of the 17 taxonomic groups, 12 guilds and 12 environments. This split Heimdallarchaeia into five guilds: 8 (Njordarchaeales), 9 (Heimdallarchaeaceae and Kariachaeaceae), 10 (Gerdarchaeales), 11 (Hodarchaeales), and 12 (Kariachaeaceae and Hodarchaeales), allowing us to determine guild-specific traits. Guild 11 includes specific enzymes like Delta3-Delta2-enoyl-CoA isomerase (ECI1, DCI), acyl-CoA:acyl-CoA alkyltransferase (OleA), olefin beta-lactone synthetase (OleC), and a gliding motility-associated binding protein (GldA). ECI is involved in auxiliary fatty acid metabolism, lipid modification, and energy production and was previously described only in bacteria and eukaryotic organelles (peroxisomes, mitochondria, glyoxysomes)^50^. Hodarchaeales in guild 11 have the entire OleABCD head-to-head hydrocarbon biosynthesis pathway, which has only been found in bacteria as a mechanism for fatty acid metabolism in lipid-rich environments^51^. Guild 11 uniquely encodes structural components known in eukaryotes for involvement in the endoplasmic reticulum and cellular organization, ORMDL^52^ and Jak and microtubule interacting protein (JAKMIP)^53^. We also discovered increased ribosomal-associated proteins within Guild 11 (Mtr4) and in Guilds 11 and 12 (L28e and Bacteroidetes VLRF1). bVLRF1 in bacteria and a few identified divergent archaea is predicted to provide additional protection from transcription and post-transcriptional processing in translation^1,54^.

We performed a pangenomic analysis on 993 single-cell eukaryotes, 437 Alphaproteobacteria MAGs, and the genomes from each of the 17 Asgardarchaeota taxonomic groups to identify orthologous groups (OGs) of genes. Asgardarchaeota and Alphaproteobacteria share 1,661 and 2,502 OGs with eukaryotes, respectively. This analysis did not include all prokaryotes, so we cannot exclude other donors, but it delineates eukaryotic OGs in each Asgardarchaeota lineage absent in Alphaproteobacteria. This identified the most functionally distinct orthologs in Lokiarchaeia, Thorarchaeia, and Heimdallarchaeia. A large number of uniquely conserved genes are present in Lokiarchaeia (1,038) and Thorarchaeia (343), which suggests their metabolic versatility as the clades with the largest number of MAGs from various environments. This genome flexibility is also reflected in the average predicted genome size (see methods) with Lokiarchaeia (Guild 1-3), Hodarchaeales (Guild 11 & 12), and Kariarchaeceae (Guild 12) guilds being greater than 4.0 Mb with a statistically significant (one-sided t-test, p value= 2.37e-16) increase in genome size from bathypelagic to epipelagic environments (Fig. 3).^55^. The five Heimdallarchaeia lineages, Hodarchaeales (215 OGs), Kariarchaeaceae (127 OGs), Gerdarchaeales (96 OGs), Heimdallarchaeaceae (83 OGs), and Njordarchaeales (39 OGs), had the next largest lineage-specific orthologs involved in a wide variety of cellular processes, including energy, amino acid metabolism, DNA maintenance, membrane biogenesis, post-translational modification, and signal transduction. This suggests that Heimdallarchaeia are more metabolically versatile than other Asgardarchaeota.

**Fig. 3.**
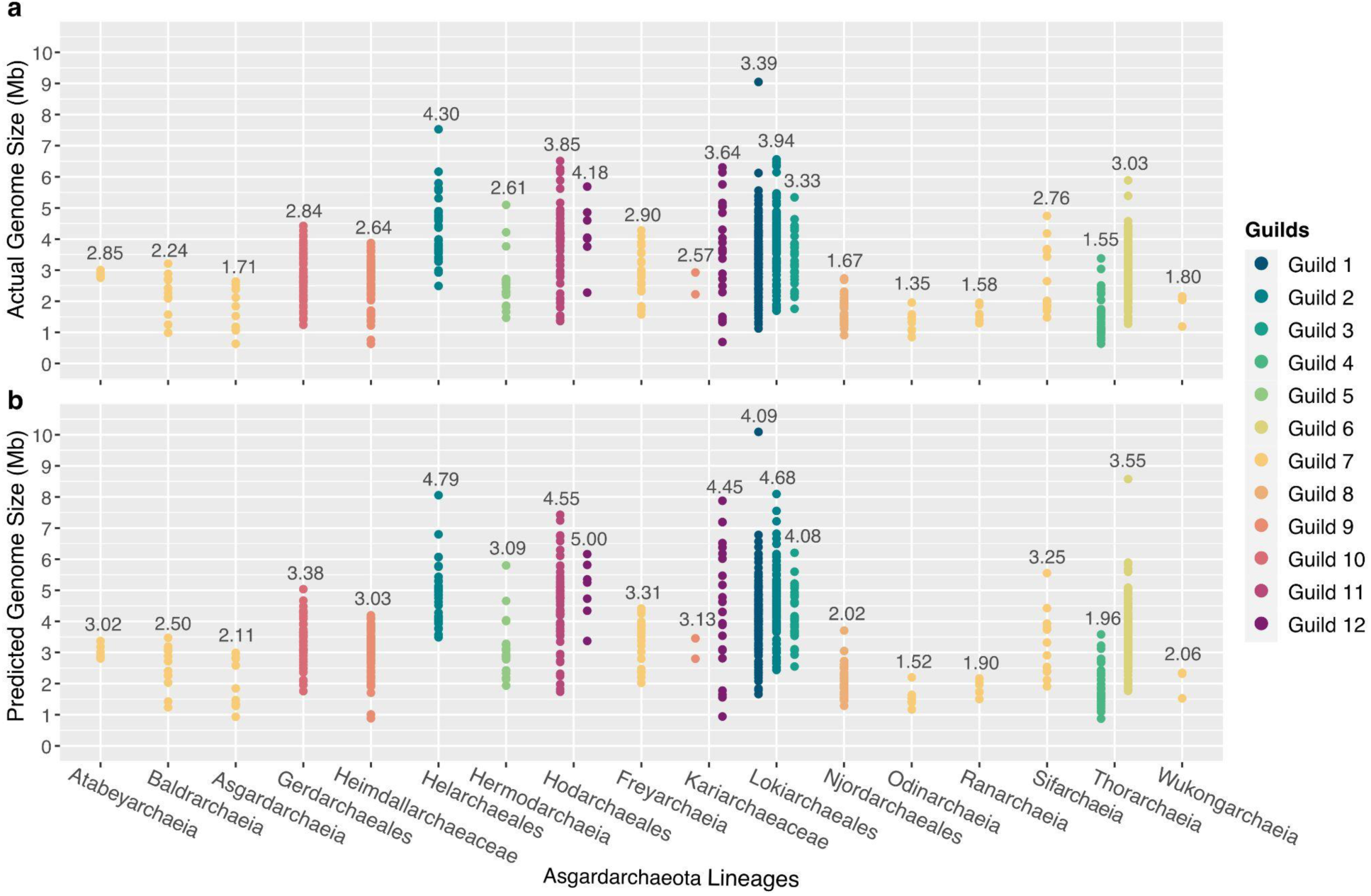
| Comparison of genome sizes of Asgardarchaeota. Actual and predicted genome size (Mb) of the Asgardarchaeaota genomic catalog separated by taxonomy and metabolic guilds. Notice members of the Heimdallarchaeia (guilds 11 and 12, Hodarchaeales and Kariarchaeaceae) and Lokiarchaeia (guilds 1 and 2, Lokiarchaeales and Helarchaeales). Listed above each subset of MAGs by lineage and guild (color-coded) is the average **a.** actual genome size (Mb) and **b.** predicted genome size (Mb) corrected for checkM v1.13 completeness and contamination (see methods).

### Novel complexes bridge the evolution of hydrogenases and Complex I

Previous studies have suggested that LAECA was hydrogen-dependent^2,56^, in contrast to contemporary organotrophic eukaryotes. To better understand the bioenergetics of this expanded Asgard archaea diversity, we profiled the distribution, structures, and evolutionarily related [NiFe]-hydrogenases and Complex I subunits (NADH:ubiquinone oxidoreductase; *nuo, E. coli* notation) across Asgard lineages^57–61^. The NuoD and NuoB subunits of Complex I are homologous to [NiFe]-hydrogenase large and small subunits, respectively^62,63^. Additionally, the membrane-bound hydrogenase (MBH) respiratory complex and Complex I share a common ancestry and structural similarity in the modular arrangement of their subunits^59,64^, forming two “arms” of the complex, a peripheral arm and a membrane arm. Both complexes couple electron transfer in their peripheral arm subunits (from ferredoxin to H_2_ and NADH to quinone, respectively) to proton pumping in the membrane arm subunits^64–66^. [NiFe]-hydrogenases were widespread among the Asgard archaea, encoded by 67% of MAGs across all 17 lineages.

We explored the potential for H_2_ exchange between the Asgard archaea and potential symbiotic partners. Phylogenetic analyses of the catalytic subunits of all [NiFe]-hydrogenases identified 18 novel subgroups, naming groups: 1m to 1n, 3e to 3l, and 4j to 4q (Fig. 4a)^42,67^. This extraordinary range of novel hydrogenases doubles the known diversity of group 3 and 4 enzymes and increases the phylogenetic resolution of the wider superfamily^67^. Structural predictions with AlphaFold2 (AF2) suggest these subunits are structurally conserved close to the catalytic center but with structurally diverse termini that may facilitate the formation of larger complexes through protein-protein interactions. 643 MAGs from 17 lineages encoded components of respiratory Complex I. We selected nine representative *nuo* gene clusters from 5 clades for structural analysis using AF2 Multimer to gain a deeper understanding of the relationships between the novel energy-converting group 4 [NiFe]-hydrogenases and Complex I.

The acquisition of new subunits and quinone reduction activity during the evolution of Complex I allowed organisms to pump additional ions, increasing the proton motive force^68,69^. AF2 Multimer predicted vast structural variation and genetic modularity of the predicted Complex I-like complexes, consistent with their phylogenetic diversity (Fig. 4a). Genomes of deep-branching Asgardarchaeota (Asgardarchaeia, Baldrarchaeia, Hermodarchaeia, and Odinarchaeia) encode unique complexes (group 4l and 4q [NiFe]-hydrogenases). These complexes share a similar subunit composition and configuration to the ferredoxin-oxidizing, H_2_-producing membrane-bound hydrogenases (MBHs) of *Thermococci*^64^. Surprisingly, they also contain a peripheral heterodisulfide reductase domain, providing a missing link between methanogenic and respiratory biochemistry^68,70^. In contrast, Heimdallarchaeia (Gerdarchaeales, Heimdallarchaeaceae, Kariarchaeaceae, Hodarchaeales, Njordarchaeales) encode novel complexes (groups 4j, 4m, 4p, 4q [NiFe]-hydrogenases) closely related to Complex I in their structural architecture. The membrane arm of these complexes is shorter than those in Complex I since they are missing one or more antiporter-like subunits. This may indicate the Heimdallarchaeia complexes transduce fewer protons than typical Complex I (but more than the Asgardarchaeia MBH complexes).

**Fig. 4.**
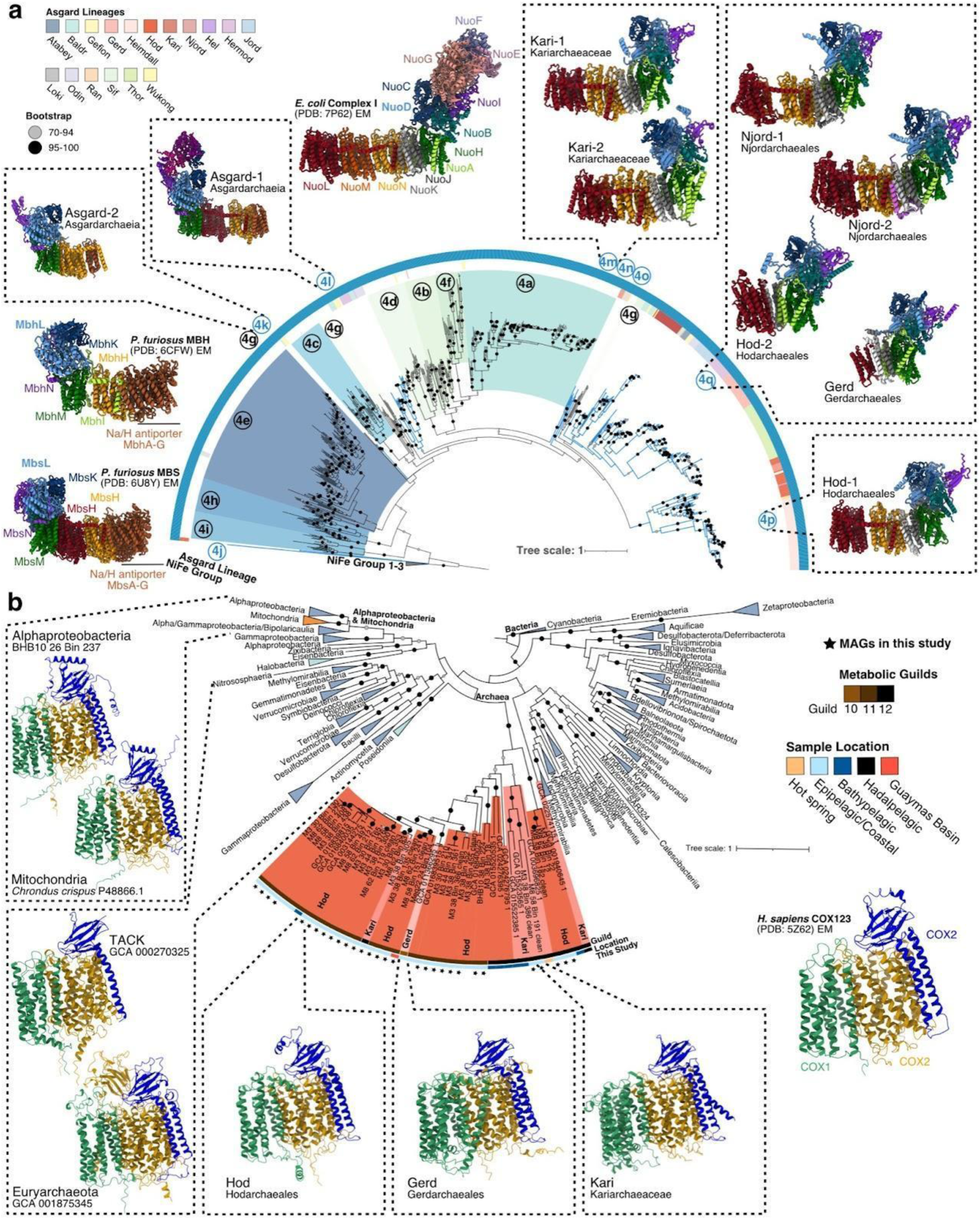
| Diversity of Complex I and IV in Asgard archaea revealing similarity in predicted structure to eukaryotes. **a.** Phylogeny of Group 4 [NiFe]-hydrogenases, displaying the novel subclades 4j-q with blue branches and Asgard lineage in the second ring. AF2 Multimer structures outline the phylogeny, indicating a transition from Mbh-like to a Complex I-like structure. Asgardarchaeia structures from 4k and 4l display an Mbh-like complex represented by EM-validated *P. furious* MBH (PDB:6CFW) and MBS (PDB:6U8Y) structures. The other complexes, 4m, 4p, and 4q, have more Complex I-like structures similar to the *E. coli* Complex I (PDB: 7P62). **b.** Maximum likelihood phylogeny based on concatenated CoxABC proteins from 343 bacteria (dark blue), 10 TACK/Euryarchaeota (light blue), 44 Asgardarchaeota (red), and 10 mitochondrial genomes (orange), rooting with cyanobacteria. The surrounding structural predictions are both CoxABC conserved structures between domains with an example cryo-EM validated structure from *H. sapiens* (PDB:5Z62). UF bootstrap values 70-94 (gray) and ≥95% (black) are shown for both phylogenies.

The peripheral arm of the Heimdallarchaeia complexes is composed of NuoBCDI subunits but is missing the NuoEFG subunits which form the NADH-oxidizing module typical of eukaryotic Complex I. Electrons may instead be provided by ferredoxin (Fd) docking to the surface of NuoI (similar to photosynthetic complex I)^71^, then transferred through a series of FeS clusters in NuoI and NuoB to the reduction site, NuoD. Asgard NuoD appears to be a catalytically active [NiFe]-hydrogenase capable of reducing protons to H_2_. This is unlike eukaryotic Complex I, which has lost hydrogenase activity for quinone reduction. Despite the prediction of large lipophilic cavities (similar to experimentally verified structures) between NuoBDH, analyses of structural models indicate that quinone is unable to enter the complex^64,72–74^. Therefore, we predict that Heimdallarchaeia Complex I-like structures reduce protons to H_2_ instead of quinone, mechanistically similar to MBH respiratory complexes, but architecturally resembling Complex I. Thus, detailed structural protein modeling using this expanded genomic catalog supports the reverse flow model, specifically, Heimdallarchaeia does not rely on a symbiont for H_2_ production^2^. Experimental studies are required to distinguish the functions of these unique complexes.

Despite the absence of NuoEFG in the Asgardarchaeota Complex I-like structures, we identified these subunits forming separate putative electron-transferring complexes. The identified NuoEFG did not form a complex with the other Nuo subunits. In Hodarchaeales, NuoEFG homologs (Complex I NADH-oxidizing module) formed a predicted complex with structural similarity to bacterial electron bifurcating and oxidoreductase complexes. The immediate up and downstream annotations indicated roles in electron transfer (FAD/NAD-binding protein and oxidoreductases) and similarity to the structurally homologous NAD-linked formate dehydrogenase. This could provide new insight into how evolution repurposes modular subunits for different electron transferring complexes, enabling NAD^+^ and Fd^+^ reduction.

The hydrogenase complexes of deep-branching Asgardarchaeia structurally resemble ancient ferredoxin-dependent hydrogenases (MBHs). Whereas, heimdallarchaeial complexes have evolved an increasingly Complex I-like structure despite the absence of quinone binding ability (quinone-dependent NADH dehydrogenases). This transition also changes the predicted FeS [4Fe-4S] stoichiometry from non-cubane (present in one subunit of MBH) to cubane (Complex I). Cubane FeS clusters with occupied coordination sites are more tolerant to reactive oxygen species (ROS), providing protection from the byproducts of aerobic respiration^75^.

### Potential for aerobic respiration in the archaeal eukaryotic ancestor

Based on the analysis of primary dehydrogenases, Heimdallarchaeia is predicted to have transitioned from the low-potential Fd-dependent metabolism of deeper-rooting Asgard archaea to reducing the higher-potential intermediate menaquinone (MK), using NADH, succinate, and possibly H_2_. Such a transition would necessitate the use of high-potential terminal electron acceptors, such as oxygen or nitrate. Nitrate reductase genes (*narGHJ*) were encoded exclusively in the deep sea and hot spring Hodarchaeales and Kariarchaeaceae, though were absent in the coastal Asgard MAGs. In contrast, the respiratory complexes required for aerobic respiration were widespread in Heimdallarchaeia (Fig. 1a). Contrary to previous reports that they are absent in Asgardarchaeota^1,2,42^, genes encoding Complex III (menaquinol-cytochrome *c* reductases; *qcrABC*) are encoded by several new MAGs from Kariarchaeaceae, Gerdarchaeales, and Hodarchaeales, as are genes for cytochrome *c* biosynthesis from heme (e.g. *ccdA*). Complex IV (cytochrome *c* oxidase; *coxABC*) is also encoded by 72 MAGs from these lineages, enabling these archaea to directly couple aerobic respiration to proton pumping.

To better understand the evolutionary history of aerobic respiration in Heimdallarchaeia, we analyzed the phylogeny, genetic organization, and structural features of Complex IV. Phylogenetic analyses of individual CoxABC proteins yielded consistent branching orders. Thus, we concatenated these proteins and this revealed they are closely related to those in the ubiquitous aerobic marine archaea Poseidoniia (Fig. 4b)^76^. These enzymes are only distantly related to those of mitochondria or their alphaproteobacterial relatives. Structural modeling analyses showed that Heimdallarchaeia Cox enzymes have similar secondary and tertiary structures compared to experimentally solved structures (RMSD of 0.82 Å between Hodarchaeales and mitochondrial structures). We found that *coxA* and *coxC* are fused in some Kariarchaeaceae MAGs. In Hodarchaeales and Kariarchaeaceae, *coxABC* genes are consistently neighbored by genes predicted to be *coxD* (predicted to assist the binding of copper ions to CoxA), the copper-binding chaperone SCO1, putatively electron-transferring DOMON-containing proteins, and the glutamine-tRNA ligase GlnRS, and were variably associated with oxidative stress and heme biosynthesis genes in a lineage, guild, and environment-specific manner. The presence of O_2_-dependent enzymes varies across Heimdallarchaeia with all lineages encoding enzymes involved in heme *o* biosynthesis, all except Njordarchaeales aromatic degradation (quercetin 2,3-dioxygenase), and only guild 12 Hodarchaeales and Kariarchaeaceae amino acid metabolism, producing substrates for the shikimate pathway (4-hydroxyphenylpyruvate dioxygenase). We found the aerobic oxidative pentose phosphate pathway is exclusive to Hodarchaeales and Kariarchaeaceae. Numerous Asgard lineages (Atabeyarchaeia, Gerdarchaeales, Heimdallarchaeia, Helarchaeales, Hermodarchaeia, Frey/Jordarchaeia, Kariarchaeaceae, Lokiarchaeales, Odinarchaeia, Thorarchaeia, and Sifarchaeia) were also capable of detoxifying reactive oxygen species (ROS), using superoxide dismutases, catalases, or peroxidases (Fig. 5). We also detected myoglobin (Mb) and protoglobin (Pgbs) homologs in 73% of the guild 12 (Hodarchaeales and Kariarchaeceae), which may have provided the ability to sense and bind oxygen at low levels.

**Fig. 5.**
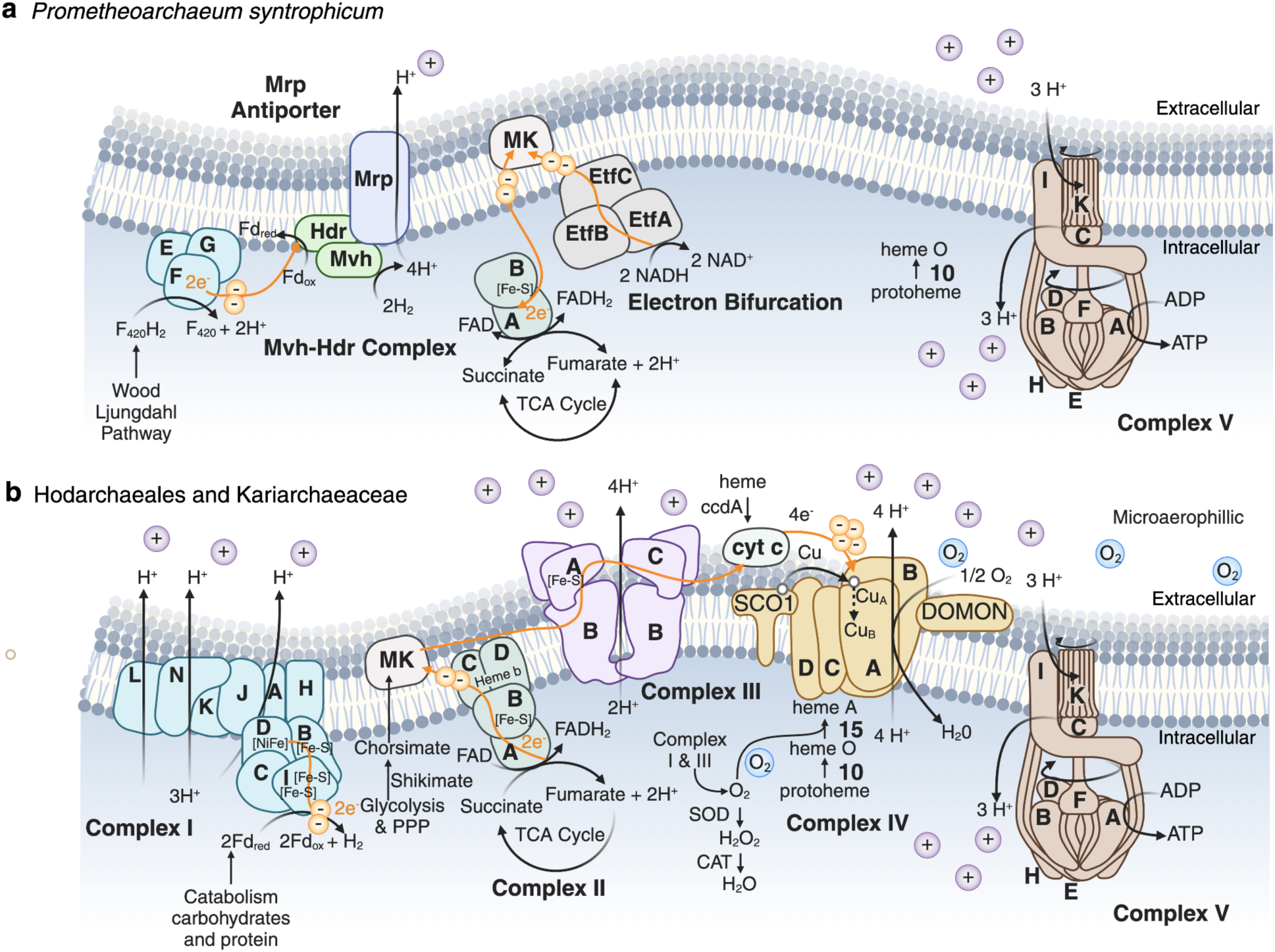
| Increasing complexity of the electron transport chain in Heimdallarchaeia compared to Lokiarchaiea. Schematic of energy conservation in **a.** the cultured Lokiarchaeales, *Prometheoarchaeum syntrophicum*, in comparison to **b.** Hodarchaealaes and Kariarchaeaceae MAGs. In the Lokiarchaeales, we identify NuoEFG components of Complex I (blue), providing electrons for the reduction of F_420_H_2_ and release of protons by the Mvh-Hdr (methyl viologen-reducing hydrogenase-heterodisulfide reductase) complex (green). The Mrp antiporter (dark purple) exports protons, providing a gradient for ATP production by V/A-type ATPase (brown; ATPVA-K). Also, there is evidence for electron bifurcation (EtfABC) and the non-membrane bound portions of Complex II (dark green; SdhAB). In contrast, we identified Complex I Nuo subunits using NADH or Fd as electron donors in Heimdallarcharchaeia. Complex I is predicted to shuttle hydrogen ions across the membrane (NuoLMN) from NADH or Fd produced in central metabolism, reducing menaquinone (MK). MK is further reduced by Complex II (SdhABCD). Electrons are passed through the Fe-S cluster in Complex III (purple; QcrABC) to cytochrome c and finally to the iron and copper metal centers in Complex IV (yellow; CoxABC4). Complex I, II, and II all help establish the proton gradient used by Complex IV (V/A-type ATPase). Additional abbreviations include O-PPP (Oxidative Pentose Phosphate Pathway) and TCA (Tricarboxylic Acid) Cycle.

### Eukaryogenesis within the context of Great Oxidation Event

The presence of a trait in a subset of modern organisms does not necessarily indicate it to be ancestral. However, aerobic quinone-dependent pathways are widespread in Heimdallarchaeia genomes obtained here and in other studies around the world. Most Hodarchaeales and Kariarchaeaceae genomes encode Complexes I (100%), II (100%), and IV (71%). Previous ancestral reconstructions predicted a transition from chemolithotrophy to heterotrophy and nitrate as the terminal electron acceptor in the Hodarchaeales ancestor^1^. We predict these organisms were dependent on organic compounds for energy and carbon sources (Fig. 6). Our findings suggest that the archaeal ancestor of eukaryotes was able to tolerate oxic conditions and utilize high-potential energy terminal electron acceptors for respiration (e.g., nitrate or oxygen). However, nitrate reduction (NarGHJ) is absent in coastal Hodarchaeales (guild 11), but present in the deep sea MAGs (guild 12). This is consistent with our phylogeny of the heme-copper oxidase superfamily, where Asgardarchaeota primarily encodes A-type heme-copper oxidases. Previous evolutionary analysis of this superfamily suggested NO reduction is not ancestral but a derived function of the ancestral A-O_2_Red^77^. This suggests early aerobic tolerance and respiration, providing essential components to remove ROS and using H_2_O_2_ for intracellular signaling. Further, vitamin B12, necessary for coenzyme synthesis, is absent in all coastal Hodarchaeales genomes. Sharing of this coenzyme has been linked to other host-symbiont metabolic interactions^78^.

**Fig. 6.**
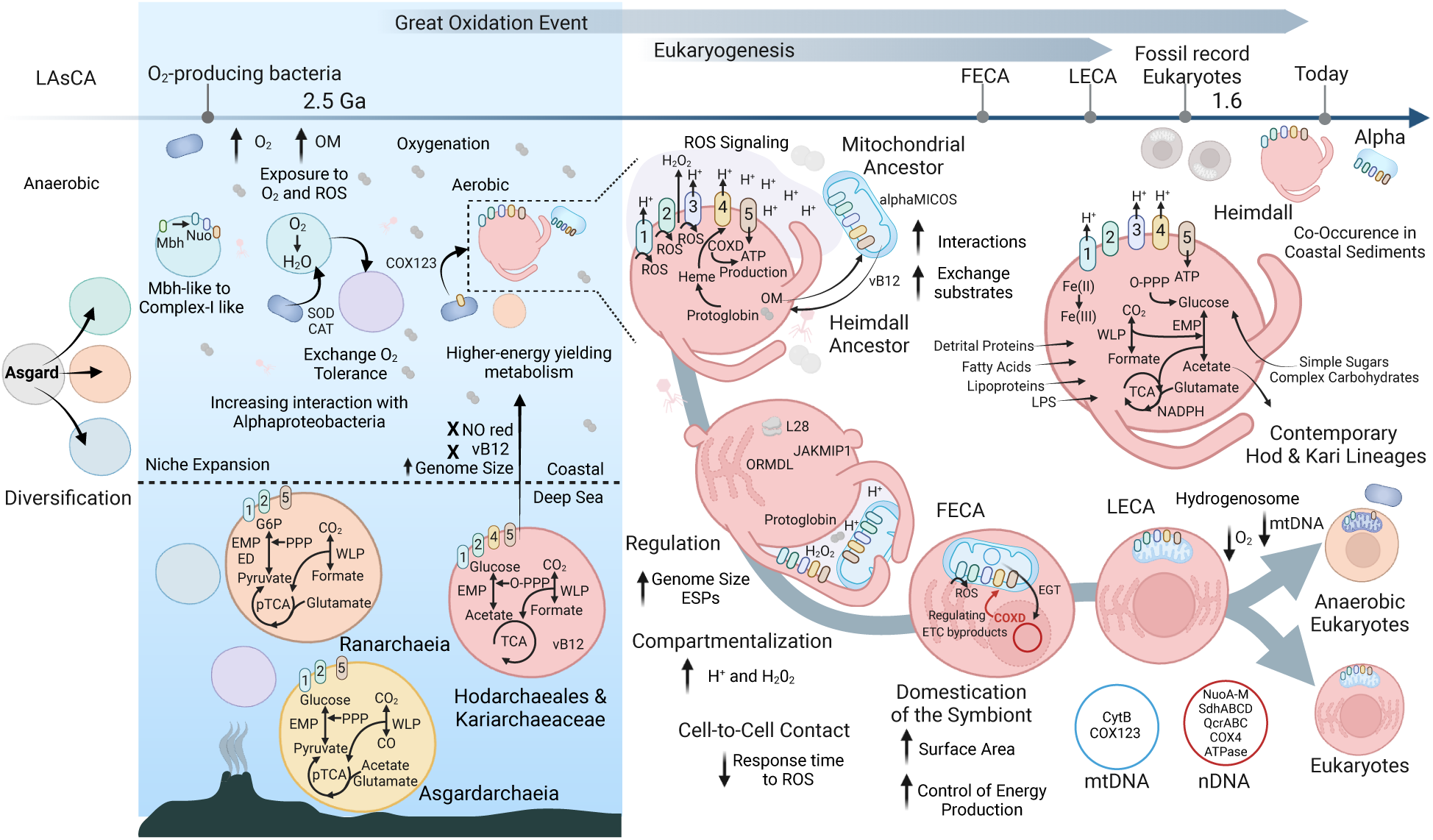
| Updated Heimdallarchaiea-centric model of eukaryogenesis in light of an expanded catalog of Asgard genomes. Our data suggest the Last Asgard archaea Common Ancestor (LAsCA) diversification allowed widespread niche expansion, including deep-sea associated lineages, Ranarchaeia (Ran; orange) and Asgardarchaeia (Asgard; yellow). Nuo subunits in several Asgardarchaeota indicate an earlier transition from MBH to Complex-I-like respiratory systems. Exposure to oxygen and the byproducts of aerobic respiration before the Great Oxidation Event triggered the exchange of oxygen tolerance and increasing interactions with the aerobic mitochondrial ancestor (blue). In Heimdallarchaeia (Heimall; red), we identify the loss of nitrate reduction (NarGHJ) and vitamin B12 (vB12) synthesis in coastal genomes present in the deep sea MAGs. The gain of ETC complexes (III and IV) increased energy yield and reactive oxygen species (ROS) production. Metabolite exchanges and coordinated response to rising oxygen levels may have increased cell-to-cell contact between partners, enabling initial compartmentalization. The Heimdallarchaeia host increased cellular regulation through ESPs (ORML, JAKMIP, L28e), using protoglobin to bind oxygen, the terminal electron acceptor. Asgard host-encoded CoxD could have controlled the endosymbiont’s ROS production in the first eukaryotic common ancestor (FECA). By the last eukaryotic common ancestor (LECA), endosymbiont gene transfer (EGT) relocated mitochondrial-encoded ETC subunits to the nuclear genome (nDNA), catalyzing the diversity of modern eukaryotes. The red arrow highlights potential regulation of ETC byproducts (ROS and ATP) by CoxD in the host’s genome. Today’s Hodarchaeales and Kariarchaeaceae are aerobic chemoheterotrophs, co-occurring with Alphaproteobacteria in oxic coastal sediments (see the main text for details). Additional abbreviations include OM (organic matter), NO red (nitrate reduction), ESP (eukaryotic signature protein), EMP (Embden–Meyerhof–Parnas Pathway), ED (Entner-Doudoroff Pathway), O-PPP (Oxidative Pentose Phosphate Pathway, NO-PPP (Non-oxidative Pentose Phosphate Pathway), pTCA (partial Tricarboxylic Acid) Cycle, and WLP (Wood-Ljungdahl Pathway).

In the Proterozoic, oxygenic photosynthesis started to change the redox chemistry of the oceans, likely fueling the transition to more complex cellular life forms capable of aerobic metabolism^79^. The timing of continental margin oxygenation is debated, but analyses indicate coastal oxic conditions ∼50 million years before the GOE^80^. An increasing abundance of Cyanobacteria in shallow water could have exposed early Asgard archaea to increasing amounts of organic matter, oxygen, and ROS. Greater cyanobacterial biomass increased the availability and preservation of organic matter, providing additional carbon sources with the greatest initial impact on shallow coastal oceans^81^. These environments are predicted to have contained the earliest eukaryotic microfossils, and now it is suggested these sediments were oxygenated at the time of fossilization^82^. Since heterotrophic metabolism is widespread among modern Asgardarchaeota, flexibility in carbon source utilization likely occurred before aerobic respiration^2^. Thus, we propose deeply rooted Asgard archaea obtained genes to survive increasing oxygenation with only some Heimdallarchaeia lineages, acquiring *coxABC* from bacteria. This provided the genetic potential to harness higher redox potentials (i.e., nitrate or oxygen) and colonize environments alongside bacteria. We identified that Heimdallarchaeia still coexist with aerobic bacteria today in various environments, across depths and sites, and with measured oxygen conditions^35–37,44^.

Heimdallarchaeia contain actin homologs are very similar to those in Lokiarchaeales, which produce protrusions, this may increase surface-to-volume ratio to access terminal electron acceptors and initial compartmentalization for concentrated proton gradients^10,18^. Portions of Complex I and III are always present in aerobic eukaryotes mtDNA. When CoxA is encoded in the nucleus it mistargets the endoplasmic reticulum (ER) instead of the mitochondrial inner membrane^83,84^. Retention of mitochondrial DNA (mtDNA) enables co-localization of redox regulation and mitigates the challenges of transferring hydrophobic subunits (i.e., mistargeting). The Inside-Out Model predicted Asgard blebs (reminiscent of now observed protrusions) gave rise to the ER, which could explain this mistargeting^85–87^. Instead of providing aerobic niche expansion, perhaps the bacterial partner(s) further partitioned metabolic pathways and ROS similar to eukaryotic peroxisomes and hydrogenosomes.

The presence of ETC components in the Asgard ancestor of eukaryotes would have provided increased energy to evolve cellular complexity, such as dynamic cytoskeleton and vesicular trafficking. If so, why would the ancestral host partner with a bacterial symbiont for aerobic respiration and then lose that capability. In light of our findings, perhaps the endosymbiont increased total respiratory capacity by expanding its membrane surface and allowed the host outer membrane to facilitate other processes, such as endo/phago/pinocytosis. This interaction could have been initially formed by the exchange of other substrates (e.g., vitamin B12). As this interaction developed, there would be redundancy in ETC in the two genomes. Our analysis supports that eukaryotic ETC components originated in bacteria. Endosymbiotic gene transfer (EGT) from the organelle to the host would have relocated symbiont subunits (e.g., NuoA-M) to the nuclear DNA. Then the Asgard ETC subunits may have been lost due to redundancy in early eukaryotes.

There is disagreement about the timing and how the host’s metabolic byproducts were shared with the bacteria that became the mitochondria. Mito-early models indicate the eukaryotic ancestor required early symbiosis to increase cellular complexity. Alternatively, mito-late models propose a highly complex host before the acquisition of proto-mitochondria and aerobic respiration^88–90^. Our analyses of extant Hodarchaeales suggests the host was less reliant on a partner(s) for energetic advantages. The added versatility in cellular processes, including ER production (ORMDL) and ribosome maintenance (L28e, JAKMIP), along with energy metabolism provide metabolic advantages to the host under appreciated in the early-late dichotomy, supporting the emergence of increased cellular complexity in mito-intermediate models^91^. In contrast, the loss of aerobic respiration in fermentative eukaryotes is instigated in low oxygen conditions, altering ETC (Complex III and IV)^92^.

## Conclusions

The generation and analyses of this expanded catalog of Asgardarchaeota advance our understanding of their contributions to the origin of the eukaryotic cell. These genomes comprise a new class, several lower taxonomic levels (49 orders and 56 families), as well as a large expansion of key metabolic protein families, including hydrogenases. This genomic dataset will be a valuable resource for further investigations into the origins of cellular complexity. In addition to providing structural components to eukaryotes, our findings indicate that Hodarchaeales also conferred bioenergetic advantages to the first eukaryotic cell. Specifically, we identified several genes predicted to code for the ETC (i.e., Complex I, III, and IV), heme biosynthesis, ROS detoxification, and protoglobins, which facilitate aerobic metabolism. Moreover, we identify Heimdallarchaeia in oxic environments and with aerobic Alphaproteobacteria (Kiloniellales, Rhodobacterales, and Sphingomonadales), suggesting interactions similar to those that resulted in the first eukaryotic cell may still be occurring. It has been proposed that regulation of the ETC arose after eukaryogenesis. However, the co-localization of CoxD and SCO1 with CoxABC in Heimdallarchaeia suggests coordination of this pathway could have originated in the Asgard archaeal host. Our analysis indicates the host and symbiont in eukaryogenesis encoded most of the ETC components. An aerobic lifestyle would have enabled the LAECA to take advantage of increasing oxygen levels during the Great Oxidation Event. This then raises the question if the archaeal ancestral host had this capability, what instigated a merger with a bacterium. Perhaps, instead of gaining aerobic respiration, the origin of eukaryotes uncoupled this process from the archaeal outer membrane to the endosymbiont (eventual organelle), increasing respiratory capacity and freeing the outer membrane to facilitate other processes, such as phagocytosis.

## Materials and Methods

### Sample acquisition

Sediment samples were collected and processed during three expeditions: BH (Bohai Sea, August 2018), GE2 (Guaymas Basin, Gulf of California, December 2008 and 2009), and GE3 (Guaymas Basin, November 2018). Bohai Sea (BH, 17-30.5 mbsl) and Guaymas Basin cores (GB, 2000 mbsl) were collected on the R/V Chuangxin Yi and HOV Alvin, respectively. Sampling through *de novo* binning from the BH and GE2 included in this study have been described previously^19,20^. Sediment samples were frozen at −80°C until extractions could take place onshore.

### DNA extraction and sequencing of Guaymas Basin Expedition 3

We split five sediment cores from 13 depth horizons from the GE3 expedition for DNA extraction. DNA was extracted using the DNeasy PowerSoil Kit (Qiagen, Germantown, Maryland, USA), quantified using a QUBIT 2.0 fluorometer (Thermo-Fisher, Singapore), and concentrated with ethanol precipitation. Extracted DNA was sent to North Carolina State Genomics Sciences Laboratory for quality control, library prep, and sequencing. Samples ran separately on an Illumina NovaSeq S4 150×2 PE flow cell (∼10 billion reads/flow cell). Three samples below the threshold for NovaSeq sequencing were sequenced with Illumina NextSeq at the University of Delaware.

We utilized FastQC and fastp v0.21.0 to assess the initial quality of the GE3 sequencing reads, removing poly-G tails (an artifact of NovaSeq sequencing), Illumina adapter sequences, reads matching human alignments, and short and low-quality reads (https://www. bioinformatics.babraham.ac.uk/projects/fastqc/)^93^. Trimmed reads were interleaved with BBTools Reformat v38.18 (http://sourceforge.net/projects/bbmap). The assembler, MEGAHIT v1.2.9, used succinct de Bruijn graphs to assemble the short-read fragments into longer contiguous reads (contigs) using a variety of kmer sizes^94^. Assemblies for each sample greater than 2.5kb were used for de novo tetranucleotide, coverage, and abundance-based binning with MaxBin v2.2.7, CONCOCT, and MetaBAT via MetaWRAP v1.3.2^95–98^. Das Tool v1.1.2-2 extracted an optimized set of non-redundant bins for each assembly from all three binning programs and MAGs were cleaned with mmgenome^99100^. The GE3 sequencing effort resulted in 5,606 archaeal and bacterial MAGs (>50% complete, <10% multiple gene copies based on CheckM v1.2.1^101^) from >10 Tb of sequencing data at 1 billion reads per sample across 13 samples, representing the largest known DNA database from the deep-sea^100^.

The reconstruction of MAGs via assembly and binning from the BH and GE2 was previously described in detail^19–21^. Briefly, the BH sequencing effort included 15 samples at 900 million reads and 120 Gb per sample, and the GE2 sequencing analyzed 16 samples at 500 billion reads and 256 Gb per sample. Resulting in a total of 5,233 BH MAGs and 3,000 GE2 MAGs >50% complete and <10% contamination based on CheckM v1.2.1^101^. Obtaining 404 Asgard MAGs required the massive sequencing effort (BH, GE2, and GE3) of 44 samples, >13,839 MAGs, and 34.5 billion reads (15 TB).

### Curation of Asgard archaea database

A comprehensive dataset was created including publicly available Asgard genomes from NCBI (as of April 2023), genomes presented in Eme et al. 2023^1^, complete genomes from wetland soils^42^, and the MAGs generated in the present study. MAGs containing gene duplications are often flagged as contamination by standard single-copy gene datasets. Before the removal of scaffolds, we compared several methods for accessing completeness and contamination (CheckM v1.2.1, miComplete v2, and MEBS v2) to avoid removing real duplicated genes which are relatively common in these archaea^101–103^. We cleaned two publicly available genomes through a combination of mmgenome and the manual removal of scaffolds due to the high copy number of single-copy marker genes^100^. MAGs with completeness >45% and contamination/redundancy below 10% based on CheckM v1.2.1 were included in the study. The resulting Asgard archaea database, ASsemblage of Genomes from Asgard Archaea Resolving DIrection to our ANcestors (ASGARDIAN), includes 140 BH MAGs, 264 GB MAGs, and 465 published MAGs.

We used this expansive genomic catalog to curate the environmental framework and metabolic annotations of these genomes. Meticulously gathered metadata from all publicly available MAGs through NCBI and publications, as well as, all collected data on the unpublished MAGs provided environmental context to this study. This data was contextualized with modeling, using amino acid frequencies to determine oxygen tolerance and optimal temperature, salinity, and pH^45^. We also used SingleM/Sandpiper 0.3.0 to search the NCBI SRA public metagenome sets prior to 2021 for OTUs linked with GTDB R220 Hodarchaeales, Kariarchaeaceae, and Alphaproteobacteria (Woodcroft et al., 2024; Eisenhofer et al., 2024).

Genome characteristics, including GC content, average amino acid identity (AAI), genome size, and abundance analyses were determined with the perl script get_gc_content.pl (https://github.com/mrmckain/), CompareM v0.1.2 with aai_wf flag (https://github.com/donovan-h-parks/CompareM), SeqKit v2.3.0, and CoverM v0.2.0-alpha7, respectively^104,105^. We conducted an initial assessment of taxonomy with Genome Taxonomy Database-Toolkit (GTDB-Tk) v2.1.1 before more extensive phylogenomic analyses^26^.

### Functional annotations

We used Prodigal v2.6.3 to predict the open reading frames in each MAG^106^. A suite of databases was used to annotate the genomes, including KofamKOALA v1.3.0 (KEGG), Interproscan v5.61-93.0-64, HADEG (Hydrocarbon Aerobic Degradation Enzymes and Genes), HydDB diamond v0.9.26.127, dbCAN2 (CAZy), METABOLIC v4.0, MEBS/Pfam v34, TIGRFAMs, EggNOGv6, and MEROPS v12.1^107–110^. Annotations were analyzed across databases and supported with gene and protein phylogenies when necessary.

### Phylogenetic analyses of the expanded Asgard genomic catalog

Four sets of markers (16S, 37 Phylosift, 47 arCOG, and NM47) were used for inferring phylogenetic relationships within this expanded diversity. We selected a representative subsample of TACK outgroup representatives, using GTDB-Tk. We chose a subset of Njordarchaeales and excluded Korarchaeia due to the hyperthermal-sensitive marker genes^1^. We extracted a subset of arCOGs (archaeal cluster of orthologous genes) with hmmsearch (v3.1b2) from 837 Asgard and 36 TACK MAGs^24^. All markers in less than 50% of the MAGs were removed from the analysis (arCOG01183). We used MAFFT v7.310 and BMGE v1.12 to align and trim the concatenated alignments. The tree was generated using RAxML v8.2.11 with free model parameters estimated by the RAxML GAMMA model of rate heterogeneity and ML estimate of alpha-parameter. Bootstraps based on 1000 rapid bootstrap inferences (Fig. 1a). Using the same set of MAGs, we compared a set of 37 single-copy marker genes from Phylosift v1.0.1^23^. The Phylosift alignment was manually accessed, masked (50% gaps) with Geneious Prime v2023.2.1, and a maximum likelihood phylogeny generated with IQ-TREE v2.0.7, using the LG+F+R10 model chosen according to BIC. We extracted the 16S rRNA genes with barrnap 0.9, using options --kingdom arc --evalue 1e-03. The length cutoff was removed to allow partial rRNA genes to be added to the analysis. To determine the geographic distribution, we blasted these against the entire Silva 138 SSUParc database with e-value 1e-05. Sequences were manually curated, aligned with MAFFT auto v7.490 and masked (50%) with Geneious Prime v2023.2.1, and ran with IQ-Tree v2.0.7^111^.

A separate analysis to test taxonomic grouping was conducted using 47 non-ribosomal proteins (here designated as NM47 for new markers), selected from a set of 200 markers previously identified as core archaeal proteins^22^, with all publicly available Asgard genomes from NCBI as of October 2022, isolated *Candidatus Lokiarchaeum ossiferum*, and Eme et al., 2023^1,18,112^. Prokka v1.14.6 was used for gene prediction for these analyses (options “--metagenome -- kingdom Archaea”)^113^. For the outgroup dataset, genus-level representatives from other archaeal lineages were selected from the Genome Taxonomy Database (GTDB), release 214^114^. Selection was based on the genome quality score (GQS), defined as GQS = completeness (%) - 5 x contamination (%), as previously described^115^. In cases where two genomes had equal GQS scores, a random selection was made between the two. The final dataset included 936 Asgard genomes and 311 genus-level representatives classified as members of the Thermoproteota (excluding Korarchaeia), Methanobacteria B, and Hadarchaeota lineages.

To recruit homologous sequences corresponding to the NM47 marker set within the selected taxa, PSI-BLAST v2.10.0+ (-evalue 1e-10) searches were conducted, using the NM47 alignments as queries against all proteomes^22,116^. All protein hits per taxon were selected and the overall sequences per marker were aligned using MAFFT L-INS-i v7.453, followed by trimming using trimAl v1.4.rev22 (-gt 0.5) and removal of sequences with more than 60% gaps^117,118^. Individual protein phylogenies were reconstructed with IQ-TREE v2.1.3 model selection from ModelFinder^119,120^. The best fitting model was selected among the combination of the LG, Q.pfam, WAG models by adding the mixture model C20 with rate heterogeneity (+R4 or +G4) with 1000 ultrafast bootstraps and calculation of SH-like approximate likelihood ratio tests (SH-aLRT)^121,122^. After removing sequences demonstrating instances of contamination, paralogy or horizontal gene transfer, the remaining sequences were realigned, trimmed and concatenated into a supermatrix containing 1244 sequences. As previously discussed in the main text, we decided to employ more complex mixture models (60 frequency vectors instead of 10) and to mitigate effects related to sequence saturation by recoding the initial untreated alignment into four character states (SR4-recoding) using BioKIT v0.0.9^123^. A phylogenetic tree was then inferred using this recoded alignment with IQ-TREE v2.1.3 under the GTR+G4+C60 model adapted to SR4 as provided by Xavier Grau-Bové (https://github.com/xgrau/recoded-mixture-models). A PMSF approximation of the selected model using the resulting tree was then used to reconstruct a final tree with 100 nonparametric bootstrap pseudoreplicates^124^. All marker sets resulted in the same taxonomic assignment for the added MAGs.

### Statistics for estimated genome sizes

To test the variability in genome size, we employed two-sided and one-sided t-tests and one-way Analysis of Variance (ANOVA) to test how actual and predicted genome size vary by environment and Asgardarchaeota taxonomic lineage. T-tests to test if water depth significantly influenced genome size and if so in what direction. For the two-sided t-test (t value = −8.3454), our null hypothesis was there is no difference in genome size for MAGs reconstructed from <1000 meters below sea level (mbsl) and >1000 mbsl. The p-value was 9.08e-15 and 4.74e-16 for actual and predicted genome size, so in both cases we rejected the null hypotheses in favor of the alternative hypotheses. Based on this analysis, there is a significant change in the actual and predicted genome sizes between the depths. One sided t-test (t value = −7.8453) allowed us to also reject the null hypothesis instead suggesting our alternative hypothesis that predicted genome size of bathypelagic and hadalpelagic MAGs are significantly (p-value=2.37e-16) smaller than epipelagic and mesopelagic. We used a one-way ANOVA with post-hoc Tukey Honestly Significant Difference (HSD) testing with an alpha value of 0.5 to check if the actual and predicted genome size is different between the isolated environments (F-value: actual= 10.32 and predicted genome size= 12.37) and taxonomic lineages F-value: actual= 21.683 and predicted genome size= 30.928). We rejected the null hypothesis that actual and predicted genome size are the same across environments and taxonomic lineages.

### Metabolic guilds definition

In order to define metabolic guilds, we determined the presence and absence profile of 19,179 protein families (Pfam v34) across the 869 Asgard MAGs with MEBSv2 (mebs.pl using the -type genomic and -comp options)^103^. Using this as the input, we utilized unsupervised clustering (mebs_clust.py) with Jaccard distance and Ward variance minimization to test various maximum distance thresholds (--cutoff) from 0.55 to 0.35 by 0.05, producing between five to 13 groups. We chose 0.35 because the 12 clusters, referred to in this study as guilds, separated the MAGs by both taxonomy and environmental parameters. Furthermore, it was at this level we could observe variance within Heimdallarchaeia. At larger distance thresholds, the few groups missed fine-scale metabolic versatility and less than 0.35 produced spurious grouping not clearly linked to taxonomic or environmental patterns, instead likely a result of genome size.

### Pangenomic analysis by lineage, guild, and environmental source

To leverage this expanded genomic catalog, we identified key transitions within the Asgard phylogeny and in comparison to the descendants of the predicted partner (Alphaproteobacteria) using pangenomics. We created presence/absence matrices for annotations from several databases (KEGG, Pfam, Eggnog) for 869 Asgard, 437 Alphaproteobacteria, and 993 single-cell eukaryotic genomes^125^. get_homologues script parse_pangenome_matrix.pl (options -m -s) was used to determine the size core, softcore, shell, and cloud of all Asgard MAG. We found 0/50/1 core, 14/372/22 softcore, 3101/8817/17382 shell, and 824/2490/9806 cloud annotations for KEGG/Pfam/Eggnog, respectively. We parsed the matrix to include results for Asgard archaea by environment, guild, and all domains by taxonomy. For the Asgardarchaeota KEGG and Pfam matrices, we used options -g to find genes/proteins present in the taxonomic group of interest (-A genomes) and absent in the other Asgard lineages (-B genomes). We did several iterations of this analysis, determining the presence of gene/protein in 5-100% of the A genomes (-P 5-100) and absent in 50-100% of the B genomes (-R 50-100).

To access functional conservation, we further parsed the Eggnog v6 output^108^. We analyzed the root orthologous group and child function to create unique identifiers for a presence/absence matrix across the domains. Matrices subset by 17 Asgard taxonomic groups, Alphaproteobacteria, and Eukaryotes were used as an input for get_homologues parse_pangenome_matrix.pl with options -m -s. Only orthologous groups conserved within the core, softcore, and shell were included, removing the cloud focused on conserved functional traits instead of recent horizontal gene transfers or *de novo* evolution less likely involved in the origin of eukaryotes. All functional OGs for Wukongarchaeia were included because there are currently too few genomes (<5) for accurate pangenome prediction by get_homologues. A list of all conserved genes was used to identify intersections across lineages and domains. Specifically, we mapped unique functional OGs shared with eukaryotes to the Asgardarchaeota branching order of the phylogeny in Fig 1b and in Eme et al., 2023^1^ to identify key transition points within the phylogeny.

### Key eukaryotic-like and metabolic genes and proteins

The pangenomic analysis highlighted several eukaryotic-like and metabolic proteins unique to taxonomy, guild, or environmental groups. To check the potential ESPs, we extracted the Asgard hits based on functional annotation and concatenated the proteins with Pfam-reviewed sequences. We used MAFFT auto v7.490, trimAl v1.4.rev15 -gt 0.5, and IQ-TREE v2.0.7 for phylogenetic reconstruction of each protein of interest. Phylogenetic reconstruction of ESPs and key metabolic proteins described in the model include GTPases, RecA, HSP90, START domain, L28e, ORML, JAKM1P1, RuBISCO, carbonic anhydrase, superoxide dismutase, catalase, protoglobin, and myoglobin. Further, MAGs encode divergent [FeFe]-[NiFe]-hybrid hydrogenases that may sense hydrogen given similarity to Group C2.

### Phylogenetic and structural analyses of the ETC subunits

To identify [NiFe]-hydrogenase groups, we ran hmmsearch v3.3.2, using hmmprofile produced by the HydDB database to extract hydrogenases from the updated Asgard genomes. The output was filtered to remove those sequences >200 amino acids and two CxxC motifs 200 residues apart (https://github.com/keappler/hydrogenase_motif_finder.py). Asgardarchaeota and HydDB reference sequences were aligned with MAFFT auto v7.490 and masked (50%) with Geneious Prime v2023.2.1. Initial alignments were accessed for additional false positives confirmed by manually accessing the alignment and AlphaFold2 (AF2) predictions of the large subunit for each novel hydrogenase clade^126^. False positives were removed and the sequences were realigned and masked, following the same methods. We reconstructed a maximum likelihood phylogeny, using IQ-TREE v2.0.7 with 1000 non-parametric bootstrap replicates (-bb 1000 -bnni) and the best-fit model, LG+R10, chosen according to Bayesian Information Criterion (BIC).

From the phylogeny we selected representatives of each novel clade for AF2 monomer predictions and the immediate upstream and downstream open reading frames (ORFs) annotated for hydrogenase or Complex I subunits for AF2 multimer predictions^127^. We note that AF2 multimer predictions may not be representative of complete complexes since ORFs may be missing from our inputs. For instance, our Mbh-like complexes are missing key Na^+^/H^+^ antiporter subunits, potentially due to our earlier functional annotation step missing these ORFs. Our AF2 predictions for [NiFe]-hydrogenase monomers and complexes were run using LocalColabFold v1.5.5^70^ (https://github.com/YoshitakaMo/LocalColabFold) installed on the Monash University MASSIVE high performance computing (HPC) cluster (www.massive.org.au). AF2 monomer predictions were run in batch with the flags --msa-mode=mmseqs2_uniref_env, --model-type=alphafold2_ptm, and --num-recycle=3 with other parameters kept on their default setting. AF2 multimer predictions were run with the flags --msa-mode=mmseqs2_uniref_env, --model-type=alphafold2_multimer_v3 and --num-recycle=48 with other parameters kept on their default setting. All predictions were computed on NVIDIA A100 Tensor Core GPUs with CUDA v12.0 installed on a MASSIVE HPC host node. AF2 outputs with the highest confidence scores (e.g. models assigned rank_001 by LocalColabFold) were used in downstream structural analysis.

We ran individual and concatenated CoxABC subunits protein trees to determine the evolutionary history of Complex IV. We added representatives from all NCBI and GTDB taxonomic orders with additional representatives from Alphaproteobacteria to 869 Asgard MAGs to determine the presence and absence in archaea and bacterial lineages. Mitochondrial genomes from RefSeq were included to place the eukaryotic subunits within prokaryotic diversity. To rule out recent lateral gene transfer events 251 Alphaproteobacteria MAGs from BH, GE2, and GE3 were included in the analysis. All references were concatenated and hmmsearchTable with hmmer v3.3 was used to extract the sequences. The sequences were filtered to only include those genomes with all three subunits, sequences were aligned with MAFFT v7.490 and trimmed with BMGE v1.12. We predicted maximum likelihood phylogeny with IQ-TREE v2.0.7 -m LG+C20+F+G and -bb 1000. To better understand the co-localization immediate upstream and downstream sequences were input into NCBI conserved domain search (CDD) and checked against other functional annotations by guild and environment for the Heimdallarchaeia MAGs encoding CoxABC.

To confirm the sequence homology of the Cox subunits A, B, and C protein complex (CoxABC), we analyzed the structural homolog of CoxABC complexes from archaeal (Asgard, TACK, Euryarchaeota), bacterial (Alphaproteobacteria) and eukaryotic organisms. Structures of the CoxABC complex acquired by cryo-EM were included in our analyses as reference structures. Protein structural modeling was performed using ColabFold v1.5.5^70^. MMseq2^128^ module and AF2^126,127^ mode were used in the modeling process combined in 2 recycling cycles. Asgard protein complex model predictions were used as queries (FoldSeek easy-search feature, >15% identity, <1E-05 e-value) to search the RCSB.org database^129^ for additional structural homologs^130^. The predicted protein structural models, the reference protein structures, and the identified structural homologues were submitted to a multi-structural alignment (MSTA) using the default parameters of mTM-align version 20220104^131^. IQ-TREE v.2.1.4-beta^132^ was used to visualize the structural homology relationship with the LG+F+R5 model chosen, according to BIC. 3D model visualization of protein complexes were done with ChimeraX software (Fig. 4b)^133^.

### Proposal of type material

Based on our described analyses, we propose the following updates to the current Asgard archaeal nomenclature, including 1 class, 49 orders, 56 families, and 178 genera. Here we provide our proposal of type material to update the nomenclature, reflecting the previous naming schemes within the recently validated Asgardarchaeota phylum^25^ and corresponding to environmental distribution of these currently uncultured clades.

We propose the naming of one new class, Ranarchaeia, and all corresponding lower taxonomic levels (Ranarchaeales, Ranarchaeaceae, Ranarchaeum, *Ranarchaeum aquilus*) within the Asgardarchaeota phylum. The consistent grouping with all phylogenetic markers used in this study and with an AAI of 40.3%-51.72% when compared to other Asgard MAGs supports this new class assignment. Because all 7 MAGs from Ranarchaeia were extracted from marine sediments, we propose to name this undescribed class “Candidatus Ranarchaeia” after the sea goddess or jötunn in Norse mythology.

*Candidatus* Ranarchaeum aquilius. *Candidatus* Ranarchaeum aquilius (aquilius. L. neut. adj. aquilius, dark). A marine-associated taxon from sediment sampling, likely chemoheterotrophs predicted to optimally prefer neutrophilic, mesohaline, and thermophilic conditions. This uncultured lineage is represented by the genome “D4993_C5_H3_Bin_486”, recovered from Guaymas Basin sediments, and defined as a high-quality metagenome-assembled genome with an estimated completeness of 89.02% and 0% contamination, 1.955 Mb genome size, 2.170 Mb predicted genome size, the presence of 2 rRNA genes (16S and 23S).

Candidatus Ranarchaeum gen. nov. (Ranarchaeum. Old Norse.fem. n. Ran, Norse goddess of the sea, N.L. neut. n. archaeum, unicellular microorganism which lack an organized nucleus; N.L. neut. n. Ranarchaeum, referring to the type genus Ranarchaeum). Type species: Candidatus Ranarchaeum aquilius.

### Descriptions of higher taxonomic ranks

Description of Candidatus Ranarchaeaceae family. nov. (Ranarchaeaceae. N.L. neut. n. Ranarchaeaceae, referring to the type genus Ranarchaeum; -aceae, ending to denote a family; N.L. fem.pl.n. Ranarchaeaceae, the Ranarchaeum family). Type genus: Candidatus Ranarchaeum.

Description of Candidatus Ranarchaeales order. nov. (Ranarchaeales. N.L. neut. n. Ranarchaeales, referring to the type genus Ranarchaeum; -ales, ending to denote an order; N.L. fem.pl.n. Ranarchaeales, the Ranarchaeum order). Type genus: Candidatus Ranarchaeum.

Description of Candidatus Ranarchaeia class. nov. (Ranarchaeia. N.L. neut. n. Ranarchaeia, referring to the type genus Ranarchaeum; -ia, ending to denote a class; N.L. fem.pl.n. Ranarchaeia, the Ranarchaeum order). Type genus: Candidatus Ranarchaeum.

## Contributions

B.J.B., K.E.A, and V.D.A. conceptualized and designed the study. B.J.B., X.G., T.J.G.E. acquired funding. B.J.B. and X.G. collected and provided environmental samples. K.E.A., V.D.A., X.G., and M.V.L. performed DNA extractions, metagenomic sequence assemblies and binning. K.E.A. collected environmental metadata and ran the functional annotation. K.E.A. and K.P. performed phylogenetic analyses. P.L. and J.P.L. ran AlphaFold2 predictions and performed structural analysis. K.E.A. conducted protein/gene phylogenetic analyses. K.E.A. and V.D.A. performed metabolic analyses. K.E.A., J.P.L., V.D.A., and B.J.B. wrote, and all authors edited and approved, the manuscript.

## Corresponding author

Correspondence to Brett Baker.

## References

1. Eme, L. et al. Inference and reconstruction of the heimdallarchaeial ancestry of eukaryotes. Nature 618, 992–999 (2023).

2. Spang, A. et al. Proposal of the reverse flow model for the origin of the eukaryotic cell based on comparative analyses of Asgard archaeal metabolism. Nature Microbiology 4, 1138– 1148 (2019).

3. Wu, F. et al. Unique mobile elements and scalable gene flow at the prokaryote-eukaryote boundary revealed by circularized Asgard archaea genomes. Nat Microbiol 7, 200–212 (2022).

4. Eme, L., Spang, A., Lombard, J., Stairs, C. W. & Ettema, T. J. G. Archaea and the origin of eukaryotes. Nat. Rev. Microbiol. 15, 711–723 (2017).

5. Spang, A. et al. Complex archaea that bridge the gap between prokaryotes and eukaryotes. Nature 521, 173–179 (2015).

6. Zaremba-Niedzwiedzka, K. et al. Asgard archaea illuminate the origin of eukaryotic cellular complexity. Nature 541, 353–358 (2017).

7. Martin, W. & Müller, M. The hydrogen hypothesis for the first eukaryote. Nature 392, 37– 41 (1998).

8. López-García, P. & Moreira, D. The Syntrophy hypothesis for the origin of eukaryotes revisited. Nat Microbiol 5, 655–667 (2020).

9. Moreira, D. & Lopez-Garcia, P. Symbiosis between methanogenic archaea and delta-proteobacteria as the origin of eukaryotes: the syntrophic hypothesis. J. Mol. Evol. 47, 517– 530 (1998).

10. Imachi, H. et al. Isolation of an archaeon at the prokaryote-eukaryote interface. Nature 577, (2020).

11. Mahendrarajah, T. A. et al. ATP synthase evolution on a cross-braced dated tree of life. Nat. Commun. 14, 7456 (2023).

12. Betts, H. C. et al. Integrated genomic and fossil evidence illuminates life’s early evolution and eukaryote origin. Nat Ecol Evol 2, 1556–1562 (2018).

13. Lyons, T. W., Reinhard, C. T. & Planavsky, N. J. The rise of oxygen in Earth’s early ocean and atmosphere. Nature 506, 307–315 (2014).

14. Lyons, T. W., Diamond, C. W., Planavsky, N. J., Reinhard, C. T. & Li, C. Oxygenation, Life, and the Planetary System during Earth’s Middle History: An Overview. Astrobiology 21, 906– 923 (2021).

15. Muñoz-Gómez, S. A. Energetics and evolution of anaerobic microbial eukaryotes. Nat Microbiol 8, 197–203 (2023).

16. Bulzu, P.-A. et al. Casting light on Asgardarchaeota metabolism in a sunlit microoxic niche. Nature Microbiology vol. 4 1129–1137 Preprint at 10.1038/s41564-019-0404-y (2019).

17. Rambo, I. M., Langwig, M. V., Leão, P., De Anda, V. & Baker, B. J. Genomes of six viruses that infect Asgard archaea from deep-sea sediments. Nat Microbiol 7, 953–961 (2022).

18. Rodrigues-Oliveira, T. et al. Actin cytoskeleton and complex cell architecture in an Asgard archaeon. Nature 613, (2023).

19. Gong, X. et al. New globally distributed bacterial phyla within the FCB superphylum. Nat. Commun. 13, 7516 (2022).

20. Langwig, M. V. et al. Large-scale protein level comparison of Deltaproteobacteria reveals cohesive metabolic groups. ISME J. 16, 307–320 (2021).

21. Gong, X. et al. Contrasting archaeal and bacterial community assembly processes and the importance of rare taxa along a depth gradient in shallow coastal sediments. Sci. Total Environ. 852, 158411 (2022).

22. Petitjean, C., Deschamps, P., López-García, P., Moreira, D. & Brochier-Armanet, C. Extending the conserved phylogenetic core of archaea disentangles the evolution of the third domain of life. Mol. Biol. Evol. 32, 1242–1254 (2015).

23. Darling, A. E. et al. PhyloSift: phylogenetic analysis of genomes and metagenomes. PeerJ 2, e243 (2014).

24. Dombrowski, N. et al. Undinarchaeota illuminate DPANN phylogeny and the impact of gene transfer on archaeal evolution. Nat. Commun. 11, 3939 (2020).

25. Tamarit, D. et al. Description of Asgardarchaeum abyssi gen. nov. spec. nov., a novel species within the class Asgardarchaeia and phylum Asgardarchaeota in accordance with the SeqCode. Syst. Appl. Microbiol. 47, 126525 (2024).

26. Chaumeil, P.-A., Mussig, A. J., Hugenholtz, P. & Parks, D. H. GTDB-Tk: a toolkit to classify genomes with the Genome Taxonomy Database. Bioinformatics (2019) doi:10.1093/bioinformatics/btz848.

27. Qu, Y.-N. et al. Panguiarchaeum symbiosum, a potential hyperthermophilic symbiont in the TACK superphylum. Cell Rep. 42, 112158 (2023).

28. Ward, L. M. et al. Geochemical and Metagenomic Characterization of Jinata Onsen, a Proterozoic-Analog Hot Spring, Reveals Novel Microbial Diversity including Iron-Tolerant Phototrophs and Thermophilic Lithotrophs. Microbes Environ. 34, 278–292 (2019).

29. Levy-Booth, D. J. et al. Genomics and metatranscriptomics of biogeochemical cycling and degradation of lignin-derived aromatic compounds in thermal swamp sediment. ISME J. 15, 879–893 (2021).

30. Seitz, K. W. et al. Asgard archaea capable of anaerobic hydrocarbon cycling. Nat. Commun. 10, 1822 (2019).

31. Speth, D. R. et al. Microbial communities of Auka hydrothermal sediments shed light on vent biogeography and the evolutionary history of thermophily. ISME J. 16, 1750–1764 (2022).

32. Reysenbach, A.-L. et al. Complex subsurface hydrothermal fluid mixing at a submarine arc volcano supports distinct and highly diverse microbial communities. Proc. Natl. Acad. Sci. U. S. A. 117, 32627–32638 (2020).

33. Zhong, Y.-W. et al. Metagenomic Features Characterized with Microbial Iron Oxidoreduction and Mineral Interaction in Southwest Indian Ridge. Microbiol Spectr 10, e0061422 (2022).

34. Liang, R. et al. Genomic reconstruction of fossil and living microorganisms in ancient Siberian permafrost. Microbiome 9, 110 (2021).

35. Rodriguez-R, L. M., Tsementzi, D., Luo, C. & Konstantinidis, K. T. Iterative subtractive binning of freshwater chronoseries metagenomes identifies over 400 novel species and their ecologic preferences. Environ. Microbiol. 22, 3394–3412 (2020).

36. Glasl, B. et al. Comparative genome-centric analysis reveals seasonal variation in the function of coral reef microbiomes. ISME J. 14, 1435–1450 (2020).

37. Rezaei Somee, M., et al. Distinct microbial community along the chronic oil pollution continuum of the Persian Gulf converge with oil spill accidents. Sci. Rep. 11, 11316 (2021).

38. Zhou, Y.-L., Mara, P., Cui, G.-J., Edgcomb, V. P. & Wang, Y. Microbiomes in the Challenger Deep slope and bottom-axis sediments. Nat. Commun. 13, 1515 (2022).

39. Bhattarai, B., Bhattacharjee, A. S., Coutinho, F. H. & Goel, R. K. Viruses and Their Interactions With Bacteria and Archaea of Hypersaline Great Salt Lake. Front. Microbiol. 12, 701414 (2021).

40. Sun, J. et al. Recoding of stop codons expands the metabolic potential of two novel Asgardarchaeota lineages. ISME Commun 1, 30 (2021).

41. Campanaro, S. et al. New insights from the biogas microbiome by comprehensive genome-resolved metagenomics of nearly 1600 species originating from multiple anaerobic digesters. Biotechnol. Biofuels 13, 25 (2020).

42. Valentin-Alvarado, L. E. et al. Asgard archaea modulate potential methanogenesis substrates in wetland soil. bioRxiv 2023.11.21.568159 (2023) doi:10.1101/2023.11.21.568159.

43. Seitz, K. W., Lazar, C. S., Hinrichs, K.-U., Teske, A. P. & Baker, B. J. Genomic reconstruction of a novel, deeply branched sediment archaeal phylum with pathways for acetogenesis and sulfur reduction. ISME J. 10, 1696–1705 (2016).

44. Tully, B. J., Graham, E. D. & Heidelberg, J. F. The reconstruction of 2,631 draft metagenome-assembled genomes from the global oceans. Sci Data 5, 170203 (2018).

45. Barnum, T. P. et al. Predicting microbial growth conditions from amino acid composition. bioRxiv 2024.03.22.586313 (2024) doi:10.1101/2024.03.22.586313.

46. Geiger, O., Sanchez-Flores, A., Padilla-Gomez, J. & Degli Esposti, M. Multiple approaches of cellular metabolism define the bacterial ancestry of mitochondria. Sci Adv 9, eadh0066 (2023).

47. Mills, D. B. et al. Eukaryogenesis and oxygen in Earth history. Nat Ecol Evol 6, 520–532 (2022).

48. Eisenhofer, R., Alberdi, A. & Woodcroft, B. J. Large-scale estimation of bacterial and archaeal DNA prevalence in metagenomes reveals biome-specific patterns. bioRxiv 2024.05.16.594470 (2024) doi:10.1101/2024.05.16.594470.

49. Woodcroft, B. J., et al. SingleM and Sandpiper: Robust microbial taxonomic profiles from metagenomic data. bioRxiv 2024.01.30.578060 (2024) doi:10.1101/2024.01.30.578060.

50. Mursula, A. M., van Aalten, D. M., Hiltunen, J. K. & Wierenga, R. K. The crystal structure of delta(3)-delta(2)-enoyl-CoA isomerase. J. Mol. Biol. 309, 845–853 (2001).

51. Rude, M. A. et al. Terminal olefin (1-alkene) biosynthesis by a novel p450 fatty acid decarboxylase from Jeotgalicoccus species. Appl. Environ. Microbiol. 77, 1718–1727 (2011).

52. Hjelmqvist, L. et al. ORMDL proteins are a conserved new family of endoplasmic reticulum membrane proteins. Genome Biol. 3, RESEARCH0027 (2002).

53. Steindler, C. et al. Jamip1 (Marlin-1) Defines a Family of Proteins Interacting with Janus Kinases and Microtubules*. J. Biol. Chem. 279, 43168–43177 (2004).

54. Verma, R. et al. Vms1 and ANKZF1 peptidyl-tRNA hydrolases release nascent chains from stalled ribosomes. Nature 557, 446–451 (2018).

55. Goyal, A. Metabolic adaptations underlying genome flexibility in prokaryotes. PLoS Genet. 14, e1007763 (2018).

56. Sousa, F. L., Neukirchen, S., Allen, J. F., Lane, N. & Martin, W. F. Lokiarchaeon is hydrogen dependent. Nat Microbiol 1, 16034 (2016).

57. Efremov, R. G. & Sazanov, L. A. The coupling mechanism of respiratory complex I - a structural and evolutionary perspective. Biochim. Biophys. Acta 1817, (2012).

58. Schut, G. J., Boyd, E. S., Peters, J. W. & Adams, M. W. W. The modular respiratory complexes involved in hydrogen and sulfur metabolism by heterotrophic hyperthermophilic archaea and their evolutionary implications. FEMS Microbiol. Rev. 37, (2013).

59. Yu, H., Schut, G. J., Haja, D. K., Adams, M. W. W. & Li, H. Evolution of complex I-like respiratory complexes. J. Biol. Chem. 296, 100740 (2021).

60. Marreiros, B. C., Batista, A. P., Duarte, A. M. S. & Pereira, M. M. A missing link between complex I and group 4 membrane-bound [NiFe] hydrogenases. Biochim. Biophys. Acta 1827, 198–209 (2013).

61. Hedderich, R. Energy-converting [NiFe] hydrogenases from archaea and extremophiles: ancestors of complex I. J. Bioenerg. Biomembr. 36, 65–75 (2004).

62. Albracht, S. P. Intimate relationships of the large and the small subunits of all nickel hydrogenases with two nuclear-encoded subunits of mitochondrial NADH: ubiquinone oxidoreductase. Biochim. Biophys. Acta 1144, 221–224 (1993).

63. Pilkington, S. J., Skehel, J. M., Gennis, R. B. & Walker, J. E. Relationship between mitochondrial NADH-ubiquinone reductase and a bacterial NAD-reducing hydrogenase. Biochemistry 30, 2166–2175 (1991).

64. Yu, H. et al. Structure of an Ancient Respiratory System. Cell 173, 1636–1649.e16 (2018).

65. Zhu, J., Vinothkumar, K. R. & Hirst, J. Structure of mammalian respiratory complex I. Nature 536, 354–358 (2016).

66. Baradaran, R., Berrisford, J. M., Minhas, G. S. & Sazanov, L. A. Crystal structure of the entire respiratory complex I. Nature 494, 443–448 (2013).

67. Greening, C. et al. Unique Minimal and Hybrid Hydrogenases are Active in Anaerobic Archaea. (2023) doi:10.2139/ssrn.4520792.

68. Schut, G. J. et al. The role of geochemistry and energetics in the evolution of modern respiratory complexes from a proton-reducing ancestor. Biochim. Biophys. Acta 1857, 958–970 (2016).

69. Chadwick, G. L., Hemp, J., Fischer, W. W. & Orphan, V. J. Convergent evolution of unusual complex I homologs with increased proton pumping capacity: energetic and ecological implications. ISME J. 12, 2668–2680 (2018).

70. Mirdita, M. et al. ColabFold: making protein folding accessible to all. Nat. Methods 19, 679– 682 (2022).

71. Schuller, J. M. et al. Structural adaptations of photosynthetic complex I enable ferredoxin-dependent electron transfer. Science 363, 257–260 (2019).

72. Kravchuk, V. et al. A universal coupling mechanism of respiratory complex I. Nature 609, 808–814 (2022).

73. Liang, Y. et al. Structure of mycobacterial respiratory complex I. Proc. Natl. Acad. Sci. U. S. A. 120, e2214949120 (2023).

74. Steinhilper, R., Höff, G., Heider, J. & Murphy, B. J. Structure of the membrane-bound formate hydrogenlyase complex from Escherichia coli. Nat. Commun. 13, 5395 (2022).

75. Bruska, M. K., Stiebritz, M. T. & Reiher, M. Analysis of differences in oxygen sensitivity of Fe-S clusters. Dalton Trans. 42, 8729–8735 (2013).

76. Rinke, C. et al. A phylogenomic and ecological analysis of the globally abundant Marine Group II archaea (Ca. Poseidoniales ord. nov.). ISME J. 13, 663–675 (2019).

77. Gribaldo, S., Talla, E. & Brochier-Armanet, C. Evolution of the haem copper oxidases superfamily: a rooting tale. Trends Biochem. Sci. 34, 375–381 (2009).

78. Jerlström-Hultqvist, J. et al. A unique symbiosome in an anaerobic single-celled eukaryote. bioRxiv 2023.03.03.530753 (2023) doi:10.1101/2023.03.03.530753.

79. Alcott, L. J., Mills, B. J. W., Bekker, A. & Poulton, S. W. Earth’s Great Oxidation Event facilitated by the rise of sedimentary phosphorus recycling. Nat. Geosci. 15, 210–215 (2022).

80. Ostrander, C. M. et al. Fully oxygenated water columns over continental shelves before the Great Oxidation Event. Nat. Geosci. 12, 186–191 (2019).

81. Canfield, D. E. Carbon cycle evolution before and after the Great Oxidation of the atmosphere. Am. J. Sci. 321, 297–331 (2021).

82. Riedman, L. A., Porter, S. M., Lechte, M. A., dos Santos, A. & Halverson, G. P. Early eukaryotic microfossils of the late Palaeoproterozoic Limbunya Group, Birrindudu Basin, northern Australia. Pap. Palaeontol. 9, (2023).

83. Roger, A. J., Muñoz-Gómez, S. A. & Kamikawa, R. The Origin and Diversification of Mitochondria. Curr. Biol. 27, R1177–R1192 (2017).

84. Butenko, A., Lukeš, J., Speijer, D. & Wideman, J. G. Mitochondrial genomes revisited: why do different lineages retain different genes? BMC Biol. 22, 15 (2024).

85. Baum, D. A. & Baum, B. An inside-out origin for the eukaryotic cell. BMC Biol. 12, 76 (2014).

86. Björkholm, P., Ernst, A. M., Hagström, E. & Andersson, S. G. E. Why mitochondria need a genome revisited. FEBS Lett. 591, 65–75 (2017).

87. Baum Buzz & Spang Anja. On the origin of the nucleus: a hypothesis. Microbiol. Mol. Biol. Rev. 87, e00186–21 (2023).

88. Pittis, A. A. & Gabaldón, T. Late acquisition of mitochondria by a host with chimaeric prokaryotic ancestry. Nature 531, 101–104 (2016).

89. Poole, A. M. & Gribaldo, S. Eukaryotic Origins: How and When Was the Mitochondrion Acquired? Cold Spring Harb. Perspect. Biol. 6, a015990 (2014).

90. Martin, W. F. et al. Late Mitochondrial Origin Is an Artifact. Genome biology and evolution vol. 9 373–379 (2017).

91. Ettema, T. J. G. Evolution: Mitochondria in the second act. Nature vol. 531 39–40 (2016).

92. Gawryluk, R. M. R. & Stairs, C. W. Diversity of electron transport chains in anaerobic protists. Biochim. Biophys. Acta Bioenerg. 1862, 148334 (2021).

93. Chen, S., Zhou, Y., Chen, Y. & Gu, J. fastp: an ultra-fast all-in-one FASTQ preprocessor. Bioinformatics 34, i884–i890 (2018).

94. Li, D., Liu, C.-M., Luo, R., Sadakane, K. & Lam, T.-W. MEGAHIT: an ultra-fast single-node solution for large and complex metagenomics assembly via succinct de Bruijn graph. Bioinformatics 31, 1674–1676 (2015).

95. Alneberg, J. et al. Binning metagenomic contigs by coverage and composition. Nat. Methods 11, 1144–1146 (2014).

96. Kang, D. D. et al. MetaBAT 2: an adaptive binning algorithm for robust and efficient genome reconstruction from metagenome assemblies. PeerJ 7, e7359 (2019).

97. Uritskiy, G. V., DiRuggiero, J. & Taylor, J. MetaWRAP—a flexible pipeline for genome-resolved metagenomic data analysis. Microbiome 6, 158 (2018).

98. Wu, Y.-W., Simmons, B. A. & Singer, S. W. MaxBin 2.0: an automated binning algorithm to recover genomes from multiple metagenomic datasets. Bioinformatics 32, 605–607 (2016).

99. Sieber, C. M. K. et al. Recovery of genomes from metagenomes via a dereplication, aggregation and scoring strategy. Nat Microbiol 3, 836–843 (2018).

100. Karst, S. M., Kirkegaard, R. H. & Albertsen, M. mmgenome: a toolbox for reproducible genome extraction from metagenomes. bioRxiv 059121 (2016) doi:10.1101/059121.

101. Parks, D. H., Imelfort, M., Skennerton, C. T., Hugenholtz, P. & Tyson, G. W. CheckM: assessing the quality of microbial genomes recovered from isolates, single cells, and metagenomes. Genome Res. 25, 1043–1055 (2015).

102. Hugoson, E., Lam, W. T. & Guy, L. miComplete: weighted quality evaluation of assembled microbial genomes. Bioinformatics 36, 936–937 (2020).

103. De Anda, V. et al. MEBS, a software platform to evaluate large (meta)genomic collections according to their metabolic machinery: unraveling the sulfur cycle. Gigascience 6, 1–17 (2017).

104. Shen, W., Le, S., Li, Y. & Hu, F. SeqKit: A Cross-Platform and Ultrafast Toolkit for FASTA/Q File Manipulation. PLoS One 11, e0163962 (2016).

105. Woodcroft, B. J. & Newell, R. CoverM: Read coverage calculator for metagenomics. Preprint at (2017).

106. Hyatt, D. et al. Prodigal: prokaryotic gene recognition and translation initiation site identification. BMC Bioinformatics 11, 119 (2010).

107. Rawlings, N. D., Barrett, A. J. & Bateman, A. MEROPS: the peptidase database. Nucleic Acids Res. 38, D227–33 (2010).

108. Hernández-Plaza, A. et al. eggNOG 6.0: enabling comparative genomics across 12 535 organisms. Nucleic Acids Res. 51, D389–D394 (2023).

109. Søndergaard, D., Pedersen, C. N. S. & Greening, C. HydDB: A web tool for hydrogenase classification and analysis. Sci. Rep. 6, 34212 (2016).

110. Hunter, S. et al. InterPro: the integrative protein signature database. Nucleic Acids Res. 37, D211–5 (2009).

111. Minh, B. Q., Trifinopoulos, J., Schrempf, D., Schmidt, H. A. & Lanfear, R. IQ-TREE version 2.0: tutorials and Manual Phylogenomic software by maximum likelihood. URL http://www.iqtree.org (2019).

112. Sayers, E. W. et al. Database resources of the National Center for Biotechnology Information. Nucleic Acids Res. 39, D38–51 (2011).

113. Seemann, T. Prokka: rapid prokaryotic genome annotation. Bioinformatics 30, 2068–2069 (2014).

114. Parks, D. H. et al. GTDB: an ongoing census of bacterial and archaeal diversity through a phylogenetically consistent, rank normalized and complete genome-based taxonomy. Nucleic Acids Res. 50, D785–D794 (2022).

115. Parks, D. H. et al. Author Correction: Recovery of nearly 8,000 metagenome-assembled genomes substantially expands the tree of life. Nat Microbiol 3, 253 (2018).

116. Schäffer, A. A. et al. Improving the accuracy of PSI-BLAST protein database searches with composition-based statistics and other refinements. Nucleic Acids Res. 29, 2994– 3005 (2001).

117. Katoh, K. & Standley, D. M. MAFFT multiple sequence alignment software version 7: improvements in performance and usability. Mol. Biol. Evol. 30, 772–780 (2013).

118. Capella-Gutiérrez, S., Silla-Martínez, J. M. & Gabaldón, T. trimAl: a tool for automated alignment trimming in large-scale phylogenetic analyses. Bioinformatics 25, 1972– 1973 (2009).

119. Minh, B. Q., et al. Corrigendum to: IQ-TREE 2: New Models and Efficient Methods for Phylogenetic Inference in the Genomic Era. Mol. Biol. Evol. 37, 2461 (2020).

120. Kalyaanamoorthy, S., Minh, B. Q., Wong, T. K. F., von Haeseler, A. & Jermiin, L. S. ModelFinder: fast model selection for accurate phylogenetic estimates. Nat. Methods 14, 587–589 (2017).

121. Hoang, D. T., Chernomor, O., von Haeseler, A., Minh, B. Q. & Vinh, L. S. UFBoot2: Improving the Ultrafast Bootstrap Approximation. Mol. Biol. Evol. 35, 518–522 (2018).

122. Guindon, S. et al. New algorithms and methods to estimate maximum-likelihood phylogenies: assessing the performance of PhyML 3.0. Syst. Biol. 59, 307–321 (2010).

123. Susko, E. & Roger, A. J. On reduced amino acid alphabets for phylogenetic inference. Mol. Biol. Evol. 24, 2139–2150 (2007).

124. Wang, H.-C., Minh, B. Q., Susko, E. & Roger, A. J. Modeling Site Heterogeneity with Posterior Mean Site Frequency Profiles Accelerates Accurate Phylogenomic Estimation. Syst. Biol. 67, 216–235 (2018).

125. Richter, D. J., et al. EukProt: A database of genome-scale predicted proteins across the diversity of eukaryotes. Peer Community Journal 2, (2022).

126. Jumper, J. et al. Highly accurate protein structure prediction with AlphaFold. Nature 596, 583–589 (2021).

127. Evans, R., et al. Protein complex prediction with AlphaFold-Multimer. bioRxiv 2021.10.04.463034 (2022) doi:10.1101/2021.10.04.463034.

128. Mirdita, M., Steinegger, M. & Söding, J. MMseqs2 desktop and local web server app for fast, interactive sequence searches. Bioinformatics 35, 2856–2858 (2019).

129. Berman, H. M. et al. The Protein Data Bank. Nucleic Acids Res. 28, 235–242 (2000).

130. van Kempen, M. et al. Fast and accurate protein structure search with Foldseek. Nat. Biotechnol. 42, 243–246 (2024).

131. Dong, R., Peng, Z., Zhang, Y. & Yang, J. mTM-align: an algorithm for fast and accurate multiple protein structure alignment. Bioinformatics 34, 1719–1725 (2018).

132. Nguyen, L.-T., Schmidt, H. A., von Haeseler, A. & Minh, B. Q. IQ-TREE: a fast and effective stochastic algorithm for estimating maximum-likelihood phylogenies. Mol. Biol. Evol. 32, 268–274 (2015).

133. Pettersen, E. F. et al. UCSF ChimeraX: Structure visualization for researchers, educators, and developers. Protein Sci. 30, 70–82 (2021).

